# Adaptation can explain cortical responses to sound deviance without requiring prediction

**DOI:** 10.64898/2026.11.14.766627

**Authors:** Przemyslaw Jarzebowski, Aditi Bishnoi, Daniel Bendor

## Abstract

Sensory systems amplify what is rare, allowing animals to orient toward threats and opportunities hidden among regular events. Whether this deviance detection reflects neuronal adaptation or error against the brain’s predictive model is debated. The two processes can contribute simultaneously because the context that enables prediction also drives adaptation. To dissociate them, we recorded from primary auditory cortex neurons in awake mice during a modified oddball paradigm that separated recent sound history from longer-term sequence statistics. Contextual effects ranged from suppression to facilitation of responses. Facilitation increased with deviance magnitude, as expected for prediction error. However, neither facilitation, suppression, nor stimulus-specific adaptation depended on these longer-term statistics. A normalization model in which a neuron’s inputs adapt at different rates reproduced the range of contextual effects and their weakening in background noise. Adaptive normalization can therefore generate deviance signals, including facilitation, without requiring a predictive model.

## INTRODUCTION

Detecting stimuli that deviate from the current sensory environment is critical for survival, allowing organisms to direct attention toward unexpected events. Deviance detection can be attributed to two mechanisms: sensory adaptation and detection of error against an internal predictive model. Both reflect a principle that sensory systems should reduce redundancy when encoding sensory stimuli and signal surprise (*1*). Both are also hard to separate experimentally because structure in the stimuli that provides predictive context simultaneously imposes adaptive pressure.

Sensory adaptation extends its effects beyond decreased spiking caused by neuronal fatigue (*2*, *3*). It tracks and adjusts the neuronal code to the statistical distribution of inputs (*4*, *5*), maximizes information transmission (*6*, *7*) and expands the dynamic range of sensory coding (*4*, *8*). Typically, preceding one sound by another has a suppressive effect (*9*) whose time course follows that of short-term synaptic depression (*10*). The suppression is stimulus-specific, as the response to a repeated sound declines while that to a rare one is spared (stimulus-specific adaptation, SSA) (*5*). Preceding sounds can also enhance neuronal firing (*11–13*) through a facilitatory effect that can be attributed to adaptation (*2*).

Enhanced responses to deviant stimuli were first characterized in EEG through mismatch nega- tivity (MMN). This event-related potential is elicited even in the absence of attention (*14*), when a rare stimulus breaks the regularity (oddball paradigm). Under predictive coding, the cortex maintains an internal model of the sensory stream and propagates the mismatch between the model’s prediction and the input (*15–17*). Whether MMN in response to deviant stimulus reflects such prediction error or adaptation has been debated (*18–20*). At the cellular level, oddball paradigms show that the spiking response to a deviant can be facilitated relative to the same stimulus when it is part of a regular pattern that is free of repetition (*21–23*). Such facilitation has been interpreted as an additive effect of prediction error (*22*, *24*, *25*), and suppression of response to the repeated stimulus as a subtractive effect of prediction (*15*, *26*). However, because oddball paradigms rely on preceding sounds to establish expectations, the resulting deviance signal is constructed in the presence of adaptation.

Separating the adaptive and predictive processes requires understanding how they interact during deviance signaling. Under adaptation, the response to a deviant depends directly on recent stimulus history and decays over time. Under predictive accounts, the deviance signal also reflects internally represented statistics of the learned sequence, so it should change with these statistics even when the immediately preceding sounds are identical (*15–17*), and should occur when the expected sound is omitted (*27*, *28*). However, global regularity violations that preserve local succession of stimuli (global oddballs) evoke weak or no deviance signals in primary sensory areas (*29*, *30*). A stronger effect of global context on deviance signals has been reported when the deviant is also a local violation (*29*), the condition in which adaptation also distinguishes the deviant from the preceding sounds. In the oddball literature, adaptation and prediction are nonetheless considered to contribute separately to deviance signals, with adaptation accounting for the suppression of repetition and prediction for the enhancement of the deviant (*22*, *24*, *25*, *31*), although adaptation alone can produce effects of either sign (*2*). We asked whether suppression and facilitation of a response to deviants arise from the same process, or whether facilitation requires predictive input.

We recorded from the primary auditory cortex of mice during a modified oddball paradigm with sound-intensity deviants. To limit accumulation of adaptation over time, we partitioned the stimulus sequence into short chunks separated by silent gaps (*29*). Within these bounded sequences, we manipulated adaptation by varying the number and timing of preceding sounds and the deviant’s physical contrast. We then tested whether global context modulates responses to local oddballs embedded in sequences with different global statistics but identical local history.

We find that context produces a continuum of single-cell response changes, from robust suppres- sion to facilitation. Both facilitated and suppressed responses increase with the sound-intensity deviance from the preceding sound, as expected for prediction error. However, facilitation, suppression, and stimulus-specific adaptation do not scale with the global statistics of the se- quence. We also find no evidence for spiking evoked by unexpected sound omissions. Instead, responses track the local stimulus history and adaptive pressure. Across the population, different rates of depression in the divisive normalization model account for heterogeneous responses to sound-intensity deviance, including response facilitation. The model also accounts for reduced facilitation and suppression when the same deviant sound is embedded in background noise. Together, these results demonstrate that adaptive normalization can generate a spectrum of contextual effects on spiking, including deviance signals, without requiring top-down predictive input.

## RESULTS

### Sound context can facilitate or suppress the spiking response

Mice passively listened to sound sequences that tested how context modulates auditory responses. The rate-level baseline sequence consisted of the same repeated harmonic chord (Fig. 1A) played at varying intensity with a 2–3 s-long interstimulus interval. This long interval minimized adaptation across the sounds (*9*, *10*). The interrupted oddball sequence additionally preceded each probe with two repetitions of the same-intensity sound (adaptor; Fig. 1B). A change in the neuronal response to the probe can be interpreted as either adaptation, because the adaptors closely preceded the probe, or prediction, because the adaptors established a regularity in the sequence that skewed the sound-intensity distribution toward the adaptor intensity.

**Fig. 1:**
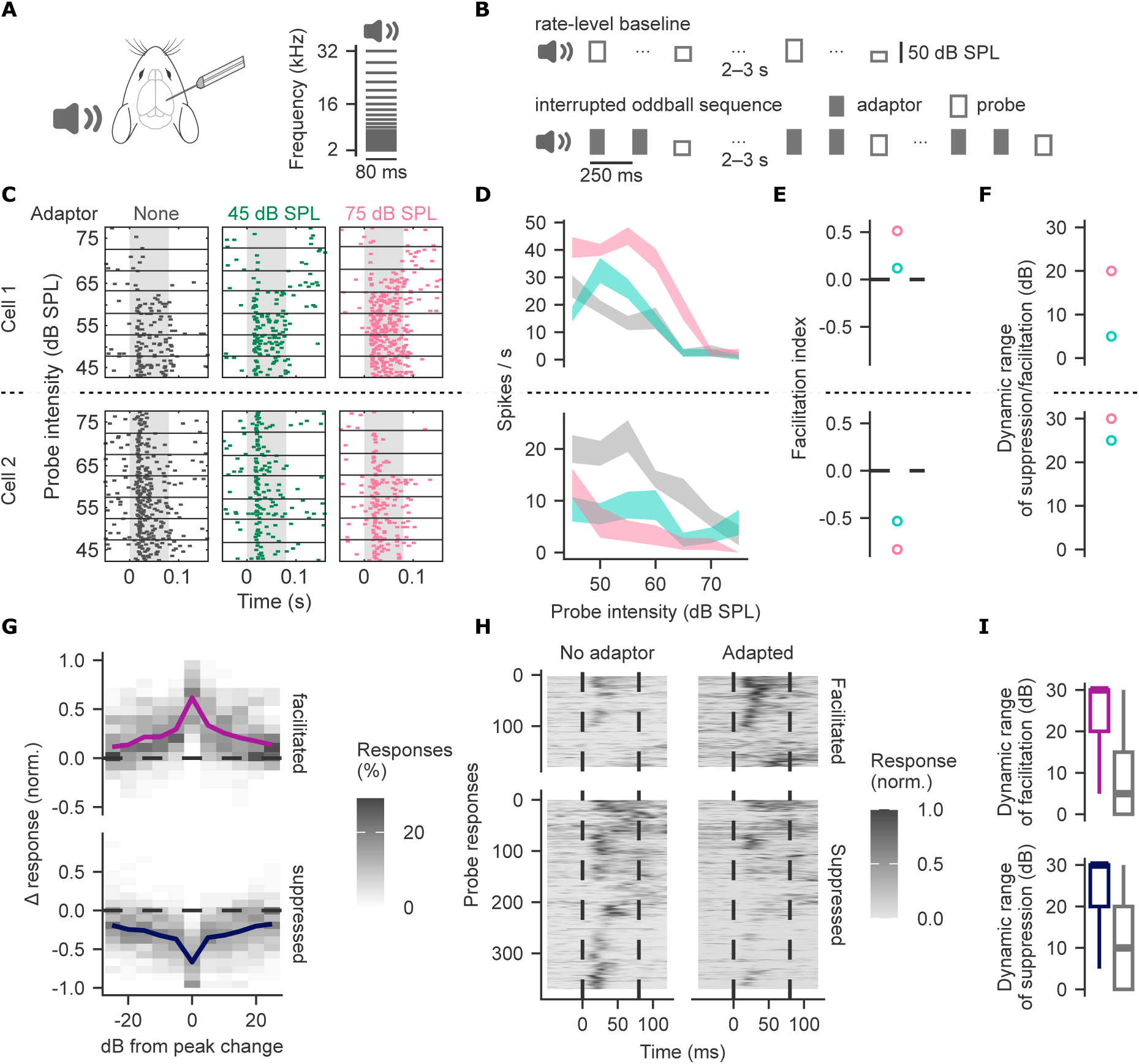
Sound context can facilitate or suppress the spiking response. **A.** Experimental setup. Awake, head-fixed mice passively listened to harmonic chords (21 log-spaced tones) presented at varying intensity. Neuropixel probes recorded from the primary auditory cortex. **B.** Stimulus paradigms. In the rate-level baseline (top), probes at 45–-75 dB sound pressure level (SPL) in 5 dB steps were separated by 2-–3 s. In the interrupted oddball sequence (bottom), the probes were preceded by a twice-repeated adaptor. **C.** Spike rasters of two example cells for probes presented in the rate-level baseline, or after a 45 or 75 dB SPL adaptor. Each row represents a single trial; horizontal lines separate probe levels. **D.** Rate-level profiles of the cells in C, without and following an adaptor. Shaded areas represent mean ± SEM. **E.** Facilitation index (FI) of each profile in D. Positive values indicate facilitation and negative values indicate suppression. **F.** Dynamic range of each profile in D: longest continuous span in the rate-level profile over which responses changed in the same direction. **G.** Change in the probe responses for all facilitated (n=186) and suppressed profiles (n=380). Profiles are aligned to the probe intensity that produced the peak change (0 dB) and are normalized to the cell’s maximum response across all probe stimuli. Greyscale shows the percentage of profiles in each bin normalized per column. Solid lines show the mean. **H.** Normalized firing rate in response to the probe intensity with the peak change in facilitated and suppressed profiles. Each row compares responses without (left) and with the adaptor (right). Dashed lines mark probe onset and offset. Responses are sorted by hierarchical clustering. **I.** Dynamic range of facilitation and suppression compared with non-significantly changed profiles (n=553). Boxes show the median and IQR; whiskers extend to 1.5 × IQR.

Of 882 units recorded in primary auditory cortex (17 sessions, 4 mice), 373 were chord-responsive (22 ± 4 per session; see Methods). Each recording session tested three adaptor intensities, yielding 1119 adapted rate-level profiles. The adaptors altered the probe responses, suppressing or facilitating parts of the profile relative to the baseline rate-level (Fig. 1C–D). We tested each profile for a change from that baseline (permutation test, p < 0.05; Methods), and assessed the change with the facilitation index (FI), defined as the modulation index of the mean adapted response relative to the mean baseline response (Fig. 1E). Profiles that changed significantly and had positive FI were classed as facilitated (186 of 1119, 16.6%), and those with negative FI as suppressed (380, 34.0%). Either change can emerge from adaptation (*9*, *11*, *12*). Alternatively, facilitation can result from the additive and suppression from the subtractive effect of prediction (*15*, *17*).

In each changed profile, we identified the probe intensity with the greatest absolute change in response. At the peak, responses increased by 61.7 ± 1.5% in facilitated profiles and decreased by 66.6 ± 1.2% in suppressed profiles, relative to the cell’s maximum response (Fig. 1G). At these probe intensities, the adaptor affected the response across the sound duration, and at the most extreme, it produced spiking in facilitated profiles where baseline elicited none, or abolished it entirely in suppressed profiles (Fig. 1H). Facilitation and suppression by the preceding adaptor generalized across a range of probe intensities: the increase spanned a continuous range of 24 ± 1 dB and the decrease 25 ± 1 dB (Fig. 1F, I, S1B). In addition to the effect on evoked responses, the adaptor increased pre-probe activity in facilitated profiles but not suppressed profiles (Fig. S1A). In all facilitated and suppressed profiles, the shifts in spontaneous rate were smaller than the absolute change in the probe response.

Different adaptors modulated the cells in a consistent direction, so a cell that was facilitated by one adaptor tended to have positive FI for other adaptors, and a cell that was suppressed by one adaptor tended to have negative FI for other adaptors (Fig. S1C). Even so, facilitation and suppression were not restricted to a specific cell population: both effects occurred across cell types (see ‘Cell type classification’ in Methods), at all cortical depths, and across cells ranging in rate-level monotonicity (Fig. S1D–F).

### Response facilitation and suppression depend on the adaptive pressure

If facilitation and suppression reflect adaptation, they should depend on the number of preceding sounds and weaken as the probe is delayed. A predictive account could produce the same dependence only if predictions emerge and decay on timescales comparable to that of adaptation.

We reduced adaptive pressure by presenting the probe after a single 75 dB sound pressure level (SPL) adaptor, and compared the probe with that at the 3rd position and in isolation (Fig. 2A). For the comparison, we only used probe intensities that were held out from the classification into facilitated and suppressed profiles (Methods), and normalized the responses to each profile’s maximum across conditions. In facilitated profiles, the probe response increased from isolation to the second position by 0.16 ± 0.05 and increased by a further 0.30 ± 0.05 at the third position (Fig. 2B–C, left). Suppressed profiles mirrored this pattern, decreasing by 0.43 ± 0.03 and then by a further 0.14 ± 0.03 (Fig. 2B–C, right).

**Fig. 2:**
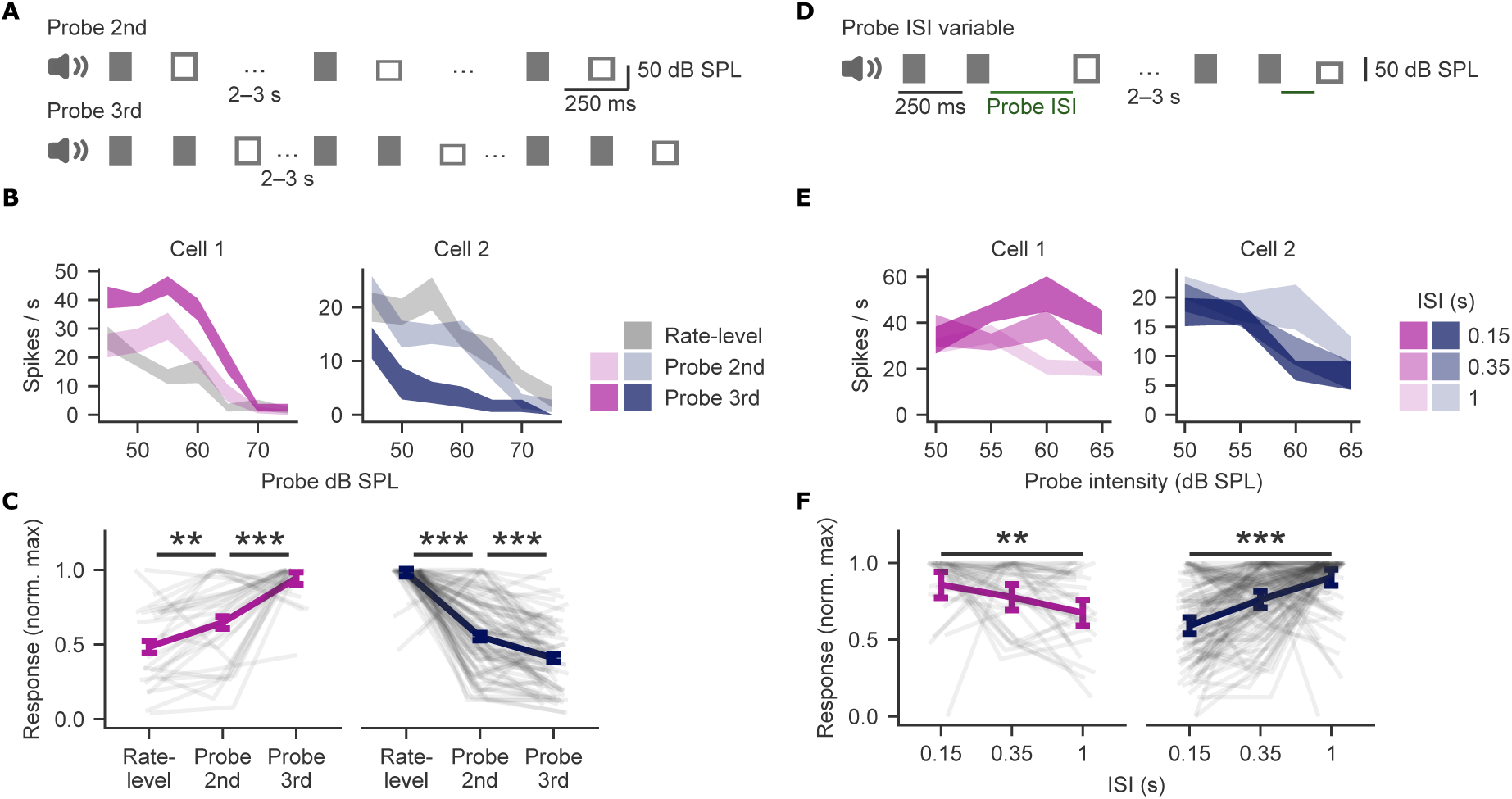
Adaptive pressure scales facilitation and suppression. A. Stimulus paradigms with the probe at the 2nd (top) or 3rd position (bottom), after one or two 75 dB SPL adaptors. **B.** Rate-level profiles of two example cells from Fig. 1 in the rate-level baseline and with the probe at the 2nd or 3rd position (mean ± SEM). **C.** Mean response of facilitated (n=29 profiles) and suppressed (n=84) by 75 dB SPL adaptor profiles plotted by probe position in the chunk. Values are normalized to each profile’s maximum across conditions. Profiles were compared on held-out levels (Methods). Grey lines show individual profiles; error bars show SEM. A linear mixed-effects model showed a position × class (facilitated vs suppressed) interaction: F_(2, 222.0)_ = 150.7, p < 10^-41^. In post-hoc pairwise comparisons, facilitated profiles increased from baseline to the 2nd position (t_(222)_ = 3.17, p = 0.005) and from the 2nd to 3rd position (t_(222)_ = 5.77, p < 10^-4^). Suppressed profiles decreased at both steps (t_(222)_ = -14.2 and -4.75, respectively; both p < 10^-4^). **D.** Stimulus paradigm with the probe presented 0.15, 0.35, or 1 s after the second 75 dB SPL adaptor. **E.** Rate-level profiles of the example cells in B at each probe ISI (mean ± SEM). Probe levels were 50–65 dB SPL. **F.** Mean response of facilitated (n=36) and suppressed (n=118) profiles by probe ISI, normalized to each profile’s maximum across the three ISIs. Grey lines and error bars as in C. The ISI × class interaction was significant: F_(2, 442.0)_ = 31.4, p < 10^-12^. Relative to 0.15 s, facilitated responses did not change significantly at 0.35 s (t_(442)_ = 1.47, p = 0.42) and decreased at 1 s (t_(442)_ = 3.32, p = 0.003). Suppressed responses increased at 0.35 s (t_(442)_ = -5.6, p < 10^-4^) and at 1 s (t_(442)_ = -10.4, p < 10^-4^). ** p < 0.01 and *** p < 0.001.

Because the number of adaptors alters sequence regularity, we next varied the probe timing (Fig. 2D), testing whether the contextual effects decay with time as expected of adaptation (*9–11*). Lengthening the interval decreased facilitation and suppression (Fig. 2E–F). Relative to 150 ms, responses in facilitated profiles did not differ significantly at 350 ms (Δ = -0.08 ± 0.05) and decreased at 1 s (Δ = -0.18 ± 0.05). Responses in suppressed profiles increased at 350 ms (Δ = 0.17 ± 0.03) and at 1 s (Δ = 0.31 ± 0.03). Both effects decayed over hundreds of milliseconds, a timescale consistent with recovery from short-term synaptic plasticity (*10*, *32*).

### Response modulation scales with sound-intensity contrast

We next investigated whether the effect of contextual modulation depends on the difference between the probe and the adaptor intensity (probe contrast; Fig. 3A). Under a predictive account, probe contrast scales the magnitude of deviance and, hence, prediction error. Under an adaptive account, the probe contrast also affects the response, because the probe response depends on how much the adaptor depresses the inputs driving it. We separately considered monotonic and non-monotonic cells (examples in Fig. S2A, Methods) because they adapt differently depending on sound-intensity statistics (*8*).

**Fig. 3:**
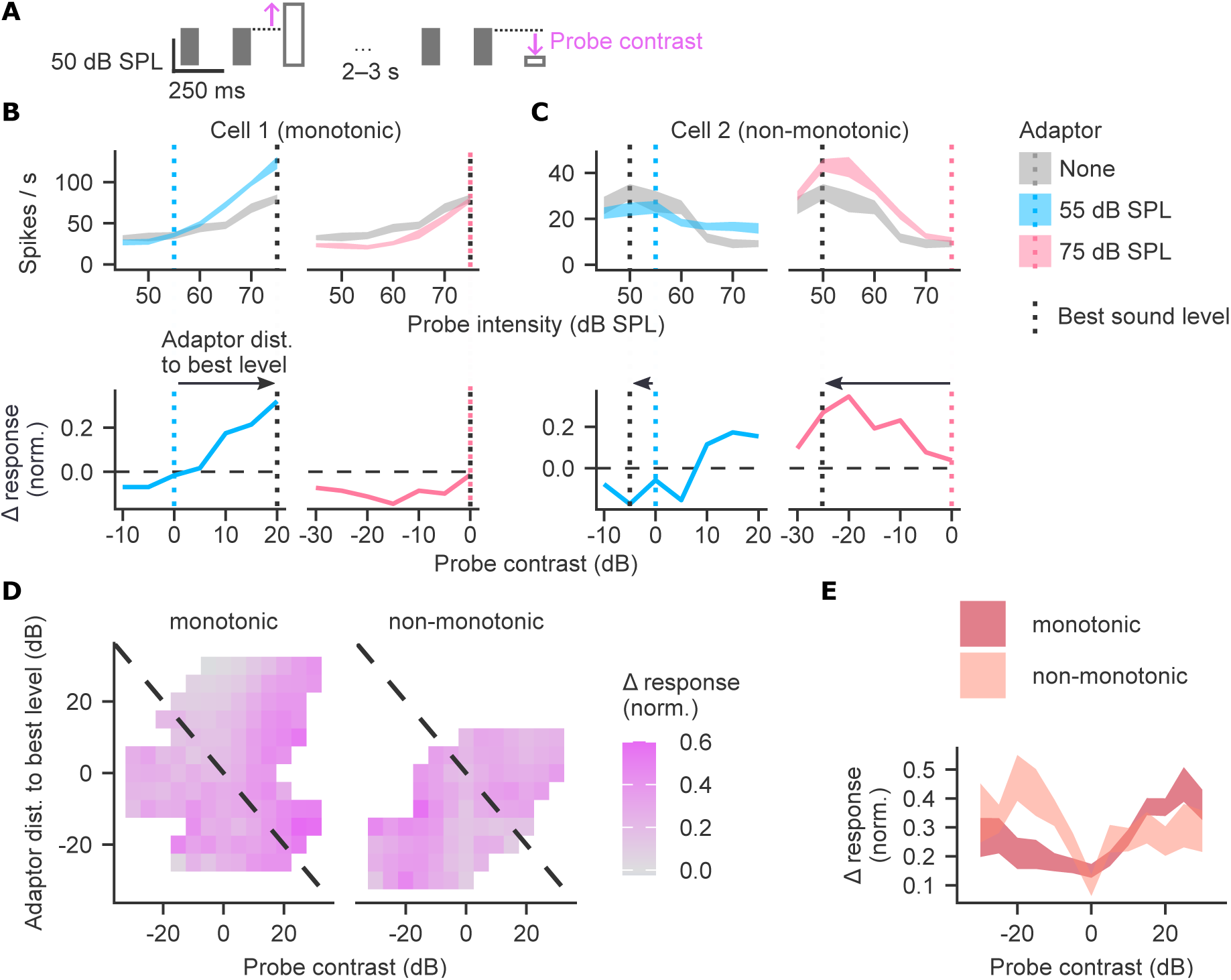
Facilitation scales with the sound-level contrast. A. Probe contrast definition. **B–C.** Example monotonic (B) and non-monotonic (C) cells with a facilitated profile. Top: responses without an adaptor and after 55 or 75 dB SPL adaptors (mean ± SEM). Bottom: change from the rate-level response as a function of probe contrast, normalized to the cell’s maximum response (across probes and conditions). Dotted lines mark the adaptor level (colored) and the cell’s best sound level (black). **D.** Mean normalized response change as a function of probe contrast and of the adaptor’s distance from the best level, for monotonic and non-monotonic cells. Data are in 5 dB bins smoothed by averaging neighboring bins; bins with samples from fewer than 5 cells are excluded. The dashed diagonal marks probes at the best level. **E.** Response change as a function of probe contrast, pooled across adaptor distances from best level (mean ± SEM).

First, we evaluated cells that had at least one facilitated profile. The cells largely preserved their monotonicity between adaptors (Fig. S2B). In monotonic cells, a low-intensity adaptor typically led to facilitation that increased with probe contrast, whereas a high-intensity adaptor slightly suppressed them (example in Fig. 3B). In non-monotonic cells, facilitation was strongest for negative probe contrasts following a high-intensity adaptor (Fig. 3C).

Statistics for D–E: a linear mixed-effects model for normalized response change (n = 1,604 responses from 231 profiles from 77 cells in 13 sessions; includes only chord-responsive cells during harmonic chords rate-level sequence). Interactions with monotonicity were significant for positive contrast (F_(1, 1590.1)_ = 7.8, p = 0.005), negative contrast (F_(1, 1557.7)_ = 14.2, p < 10^-3^) and adaptor distance (F_(1, 617.9)_ = 6.0, p = 0.01). Slopes per dB with 95% CIs are given in the Results.

Facilitation depended not only on the probe contrast but also on which adaptor preceded the probe (Fig. S1C, Fig. 3B–C). We therefore described the contextual modulation as a joint function of probe contrast and the adaptor’s distance from the cell’s best sound level (Fig. 3D). The two coordinates sum to the probe’s level relative to the best level. Averaged across adaptors, facilitation increased toward both extremes of probe contrast (Fig. 3E). We therefore modeled the response change with positive and negative contrast as separate predictors (Methods). This described the data better than a model with a single probe contrast term (χ²(1) = 71.3, p < 10^-15^; ΔAIC = −69.3).

The monotonicity groups differed significantly in how probe contrast sign scaled facilitation (Fig. 3D–E). In monotonic cells, facilitation increased with positive probe contrast (95% CI [0.01, 0.013]), but not with negative contrast (95% CI [-0.003, 0.001]). In non-monotonic cells, it increased with both negative and positive probe contrast (negative: 95% CI [0.004, 0.009]; positive [0.002, 0.008] per dB). The effect of adaptor distance also differed between monotonicity groups. In monotonic cells, facilitation increased as the adaptor approached and exceeded the best level (95% CI [0.0003, 0.0031] per dB). Non-monotonic cells showed no significant change (95% CI [−0.0037, 0.0007] per dB).

Next, we tested how deviance magnitude scales suppression in cells with at least one suppressed profile. An analogous model of suppression magnitude showed that suppression was strongest near zero probe contrast and weakened as contrast increased. This held in both directions (positive and negative contrast) in monotonic cells and for positive contrast in non-monotonic cells (Fig. S2C–F).

In both facilitated and suppressed cells, the response relative to the rate-level baseline scaled with the deviation from the adaptor: facilitation increased while suppression decreased. This scaling with deviance magnitude is consistent with prediction error (*15*, *17*), but it does not by itself separate prediction from adaptation. Therefore, we next manipulated sequence statistics and regularity while leaving the local stimulus history unchanged.

### Deviance signals depend on local stimulus history, not on global sequence statistics

#### Response facilitation and suppression do not scale with sequence statistics when local history is matched

Adaptation depends on recent stimulus history alone, whereas a predictive model also tracks sequence statistics. Because two sequences with different statistics would lead an ideal observer to form different predictive models, the prediction error should differ between them. To investi- gate whether prediction contributes to deviance signals, we compared responses between two stimulus blocks with local deviants that had identical local history but different global statistics (Fig. 4A).

**Fig. 4:**
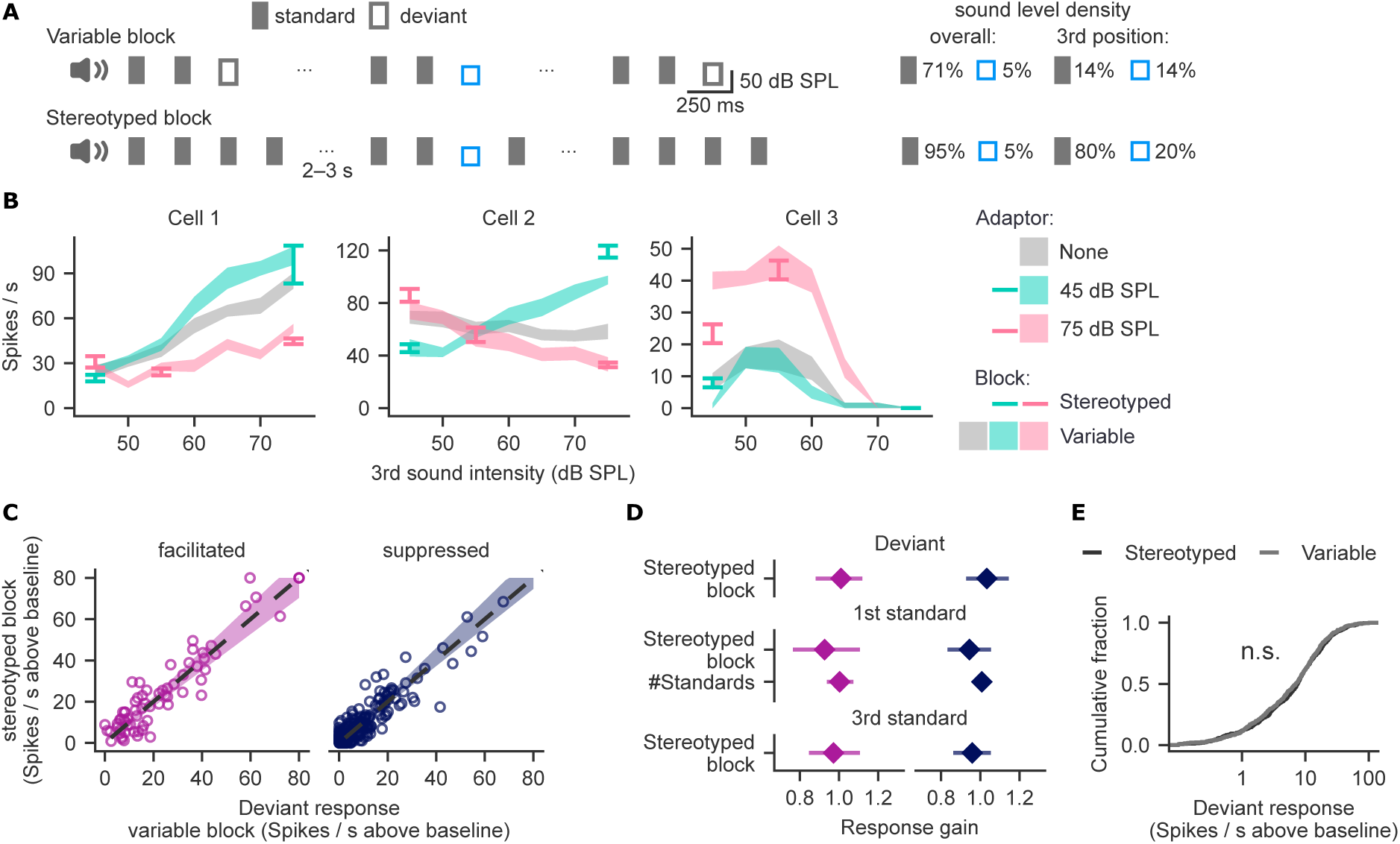
Response facilitation and suppression do not scale with sequence statistics when local history is matched. A. Variable and stereotyped blocks (left), with matched local history but different global statistics (right). **B.** Responses of three example cells to deviants in the variable (bands) and stereotyped blocks (error bars), shown as mean ± SEM. We tested three stereotyped blocks: two with a 75 dB standard and different deviants, and one with a lower-intensity standard and a higher-intensity deviant. **C.** Deviant responses in the stereotyped versus the variable block. Profiles were compared on held-out levels (Methods). The dashed line marks equality; the colored band shows the 95% HDI of the gain from a Bayesian hierarchical model. Data for deviant responses significantly exceeding spontaneous activity in either block, facilitated responses: n = 69 from 58 profiles from 36 cells; suppressed responses: n = 222 from 173 profiles from 91 cells. Poisson mixed model of spike counts: facilitated, χ²(1) = 0.12, p = 0.73; suppressed, χ²(1) = 0.001, p = 0.98. **D.** Posterior densities of the response gain in the stereotyped relative to the variable block. For facilitated deviant responses: median 1.01, 95% HDI [0.89, 1.12]; for suppressed: 1.03, [0.93, 1.14] (data as in C). For facilitated first standard: 0.93, [0.76, 1.10]; suppressed: 0.94, [0.84, 1.05]; gain per additional standard in the preceding chunk: 1.00, [0.93, 1.08] and 1.01, [0.98, 1.04] (data as in S3A). Third standard: 0.98, [0.85, 1.12] and 0.96, [0.87, 1.06] (data as in S3C). **E.** Cumulative distributions of deviant responses in the two blocks. Data for deviant responses significantly exceeding spontaneous activity in either block (n = 656 responses per block from 508 profiles and 200 cells). Kolmogorov–Smirnov test, p = 0.77.

The first block was the interrupted oddball sequence. In it, the third sound took one of seven intensities with equal probability, and so typically differed from the preceding two (variable block; 6/7 probability of sound-intensity change). In the second block (stereotyped block), each chunk repeated the same intensity sound four times. In 20% of chunks, the third position instead had a deviant intensity that was fixed across the block. The deviant was a local change in both blocks; it was expected in the variable block but violated a global pattern in the stereotyped block. Mice were exposed to both variable and stereotyped blocks during the habituation period over 3 days.

The predictive models we considered make different predictions for the two blocks (Table S1). A predictive model that tracks the global pattern entails a larger prediction error for deviants in the stereotyped block. Models that track the probability of each sound intensity predict a block difference for either the deviant or for standard, and the direction of the difference depends on how the model represents the expected stimuli.

We compared deviant responses between the two blocks, taking profiles and probe intensities at which the response was facilitated or suppressed in the variable block (Fig. 4B–C). We modeled the response change with a Bayesian hierarchical model of spike counts (Methods). The model estimates a gain: the factor by which the response in the stereotyped block differs from that in the variable block (gain of 1 means no change). Gains were close to 1 for facilitated responses (95% HDI [0.89, 1.12]) and suppressed responses (95% HDI [0.93, 1.14]; Fig. 4D). Individual profiles could still deviate from the group gain, but no facilitated profile did (Methods). Having found no difference for the deviant between variable and stereotyped blocks, we compared responses to the standard, whose density rises from 71% in the variable to 95% in the stereotyped block. We first took the standard at the first position, which follows 2–3 s of silence, and included the number of standard repetitions in the preceding chunk as a covariate to account for adaptation across chunks. The covariate had no effect, and the first-standard response did not differ between blocks (Fig. 4D, S3A). A model of the response to the third standard gave similar evidence for no change (Fig. 4D, S3C). The lack of change in the deviant and the standard does not support any of the predictive models in Table S1.

The analyses above assumed that prediction error scales the facilitated and suppressed responses in the same direction in the variable and stereotyped block. To test for changes that do not follow this assumption but still scale overall population responses, we compared distributions of deviant and standard response magnitudes across chord-responsive cells. Neither differed between the blocks (Kolmogorov-Smirnov test; Fig. 4E, S3B, D).

#### Stimulus-specific adaptation does not change with global sequence statistics

So far, we assessed contextual effects relative to the sound presented in isolation. A complemen- tary approach measures stimulus-specific adaptation (SSA) by comparing responses to the same sound presented as a standard and as a deviant. Some of that relative increase in the deviant response has been proposed to reflect prediction error (*22*, *24*), which should depend on global sequence statistics. We compared responses in two symmetric configurations of the sound stimuli: one in which the standard was 50–55 dB SPL and the deviant was 75 dB SPL, and one in which the roles were reversed. We combined the two into the common SSA index (CSI; Fig. S4A) (*33*). In the variable block, the median CSI was positive (Fig. S4B–C), indicating that adaptation was specific to sound intensity, consistent with previous reports on intensity deviants (*5*; but see *34*).

Next, we investigated whether SSA scales with the global sequence statistics. The CSI distri- bution across the neuronal population did not differ between variable and stereotyped blocks (Fig. S4D–E). The hierarchical Bayesian model estimated the shift in the CSI as negligible (Fig. S4F). SSA of intensity deviants was therefore invariant to global sequence statistics.

#### Unexpected sound omission does not evoke a spiking response

In some predictive coding models, omitting an expected sound generates spiking because the prediction does not match the sensory input (*27*, *28*). We tested this in a separate cohort of mice that were exposed only to stereotyped sequences of five identical sounds without deviants during habituation. Their expectation of each sound should therefore be at least as strong as in the main cohort. During recording, the third sound was omitted in 20% of trials (Fig. S5A).

None of the 201 cells recorded in the primary auditory cortex increased their spiking above a z-score of 1 during the 80 ms window aligned with the omitted sound (n=3 animals; 11 sessions; Fig. S5B–C). No cell significantly changed its activity during the omission period relative to the immediately preceding period (Wilcoxon signed-rank test, p < 0.05). Finally, activity during the omission did not deviate strongly from that in a post-sequence control window, which is subject to comparable post-stimulus rebound activity but not to any deviance signals (Fig. S5D).

Thus, auditory cortex spiking responses neither scaled notably with global sequence statistics nor increased during the omission of an expected sound.

### Continuum of adaptation rates explains heterogeneous responses to deviant sound intensity

Because facilitation and suppression showed no properties beyond those explained by adaptation, we sought a computational model that produces both from a single adaptive process. Within individual profiles, the adaptor’s effect varied with probe intensity (Fig. 5A–B), so it could not be described as a fixed additive or multiplicative effect. Across the population, the effects of adaptation spanned a continuum from suppression through no change to facilitation, without distinct boundaries (Fig. 5C). This continuum argues for a graded mechanism of adaptation.

**Fig. 5:**
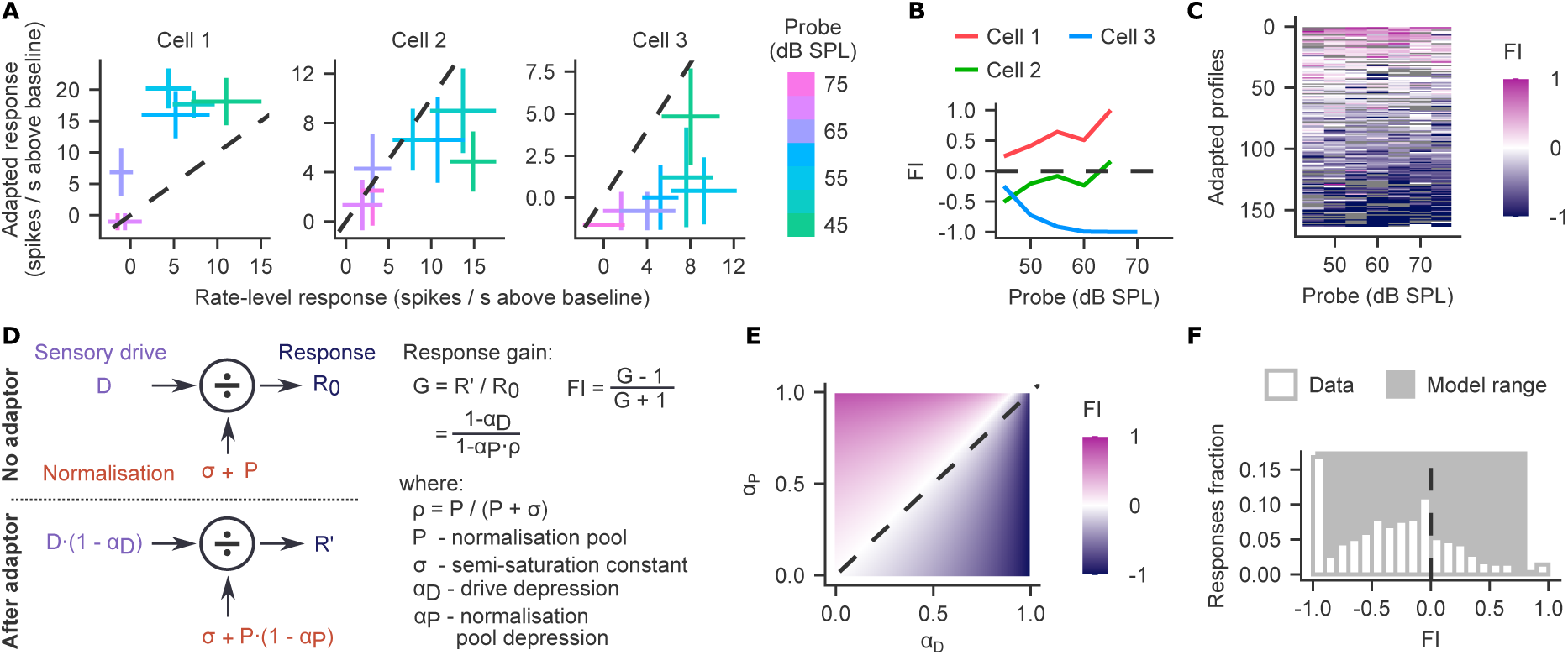
Continuum of adaptation rates explains heterogeneous responses to deviant sound levels. A. Adapted versus rate-level responses at each probe level for three example cells after a 75 dB SPL adaptor: facilitated (Cell 1), not significantly changed (Cell 2) and suppressed (Cell 3). Crosses show mean ± SEM; dashed lines mark equal responses. **B.** FI of the responses in A as a function of probe level. FI is shown only where driven the sum of the rate-level and adapted responses exceeded twice their SEM. **C.** FI at each probe level for profiles adapted with 75 dB SPL, sorted by mean FI. Only profiles of cells whose mean driven firing rate in the rate-level baseline exceeded 2 Hz are shown (n=163). Grey marks probe levels excluded by the criterion in B. **D.** Divisive normalization model without (top) and with (bottom) an adaptor, which depresses the sensory drive and the normalization pool. **E.** FI predicted by the model as a function of drive and pool depression. The dashed line marks equal depression. **F.** Distribution of FI across all probe responses in C. The dashed line marks FI = 0. The histogram peak at -1 corresponds to the adaptor fully suppressing spikes, and at 1 to the adaptor facilitating a previously unresponsive cell.

Synaptic depression has been proposed to be the mechanism of short-term adaptation in the auditory cortex on timescales that can span the interstimulus interval between the adaptor and probe (*10*). Because synaptic depression can only weaken neuronal inputs, a model in which depression acts on the excitatory drive alone cannot produce facilitation. We previously showed that depression of excitatory and inhibitory inputs can increase and decrease auditory spiking across a stimulus train (*35*). Inhibitory interneurons form a pool that can normalize the sensory- driven response. Divisive normalization model formalizes this computation by placing this normalization pool in the denominator, where it sets response gain (*36*, *37*). Depression of excitation and of inhibition becomes depression of the sensory drive and of the normalization pool, where the latter can enhance the response (*2*). The resulting response gain depends on these two depression rates (Fig. 5D).

To validate the model, we simulated the gain across combinations of the two depression rates (Fig. 5E). Facilitation occurs when the normalization pool is sufficiently more depressed than the drive; otherwise, the response is suppressed (Fig. 5E). Balanced depression of both pathways yields weak suppression, a common effect in our data (Fig. 5C, F). The model, parametrized by independently varying these two rates, generated the full range of suppression we observed and most of the facilitation (Fig. 5F). The model’s facilitation is bounded because ρ remains affects the response even when the pool is fully depressed (Methods). By contrast, the observed FI saturates at ±1 whenever either the baseline or adapted response evoked no spikes.

### Background noise reduces the adaptor’s effect

If facilitation and suppression arise from depression of the sensory drive and the normalization pool, they should weaken when the adaptor engages these two pathways less effectively. We tested this by presenting the adapting sequences in noise (Fig. 6A), which reduces sound-evoked responses (*38–40*). Of the 373 chord-responsive cells, 209 were also chord-responsive in low or high noise and were included in the comparison.

**Fig. 6:**
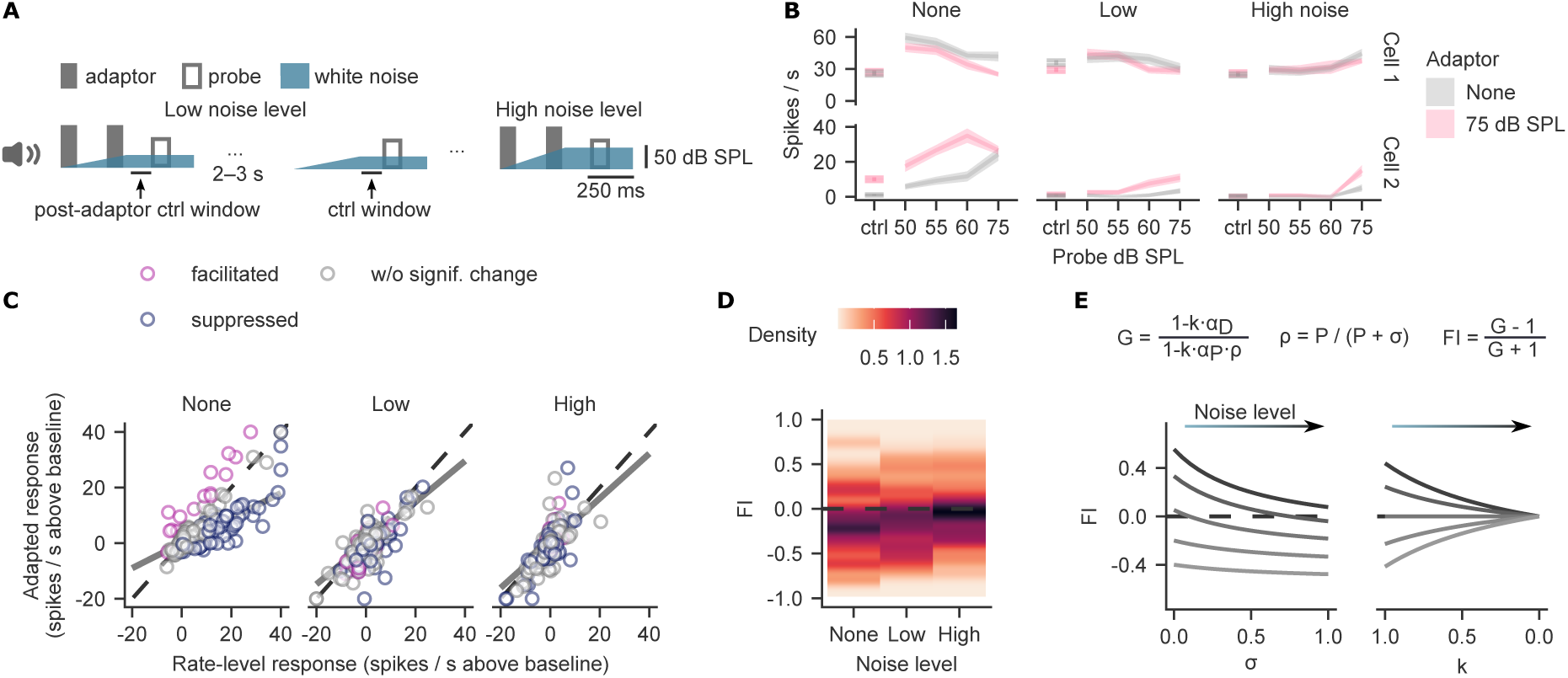
Background noise reduces the adaptor’s efficacy in depressing the sensory drive and normalization pool. **A.** The interrupted oddball sequence (Fig. 1) was embedded in a low or high level of white noise. Noise ramped up from the onset of the first adaptor, reaching full amplitude at the offset of the second adaptor (adaptors at 75 dB SPL). In the rate-level control, probes were embedded in the same ramping noise without adaptors. Horizontal bars mark the pre-probe windows compared in B. **B.** Rate-level profiles of three example cells without noise and in low and high noise, without or with an adaptor (mean ± SEM). Rate-level profiles include four probe intensities tested in noise. ”ctrl” shows activity in the control window. **C.** Mean adapted versus mean rate-level baseline response of each profile plotted at different noise levels. Profiles are colored by their classification in the noise-free condition. The dashed line marks equal responses; the solid lines mark the estimated response gain. Bayesian hierarchical model of the adaptor’s gain, no noise: median 0.45, 95% HDI [0.36, 0.55]; low noise: 0.74, [0.56, 0.92]; high noise: 0.80, [0.46, 1.18]. **D.** FI across profiles at each noise level, shown as density. The dashed line marks FI = 0. **E.** Simulated effect of noise on FI for example combinations of α_D_ and α_P_ (see Methods). Left: noise modeled as an increase in σ, with k = 1. Right: noise modeled as a decrease in the adaptor efficacy k. The x-axis is reversed to show the effect of increasing noise level from left to right.

Noise increased firing in the control window preceding the probe and reduced evoked responses (Fig. S6A–B), consistent with previous reports that background noise reduces and shifts rate- level functions (*38–40*). Before comparing evoked responses, we assessed how the noise-driven activity in the control window changed following the adaptor. It did not significantly change in facilitated profiles (Fig. S6C, left), arguing against nonspecific disinhibition as an explanation for facilitation. Noise-driven activity decreased slightly in suppressed profiles (Fig. S6C, right).

Evoked responses changed in opposite directions: the previously facilitated profiles scaled down and the suppressed profiles scaled up, both converging on the rate-level response in noise (Fig. 6B–C). We quantified the adaptor’s effect on driven responses relative to the rate-level baseline using a Bayesian model that estimated gain separately at each noise intensity. Across all profiles, the gain rose with noise (Fig. 6C). Consistent with convergence of adapted responses to the rate-level baseline, the fraction of profiles with |FI| ≤ 0.2 increased from 35% without noise to 49% in high noise (bootstrap 95% CI: [4%, 23%]). The FI interquartile range narrowed non-significantly from 0.48 to 0.36 (95% CI: [-0.18, 0.05]; Fig. 6D).

We then considered what change in the normalization model accounts for this convergence. Noise could increase the non-depressing part of the normalization signal, equivalent to raising *σ*. Raising *σ* lowers the overall gain, causing the facilitation to disappear, but it also reduces gain in suppressed profiles (Fig. 6E, left), contrary to our data. Alternatively, noise could reduce the adaptor’s efficacy *k*, a common factor scaling α_D_ and α_P_. This change could reflect lower evoked responses to the adaptor, leading to less depression of both pathways, or noise-driven activity already depressing both pathways. Reducing the efficacy *k* diminished facilitation and suppression, leading to convergence of adapted responses towards the baseline as observed in the data (Fig. 6E, right).

## DISCUSSION

Because sensory history shapes adaptation and prediction, these two processes are usually entangled. We examined how they shape deviance signals in the primary auditory cortex. A short, predictable sequence strongly modulated neuronal responses to a subsequent deviant sound. The effects ranged from suppression to facilitation relative to the same sound presented in isolation (*11*, *12*). Consistent with adaptive accounts of deviance, facilitation and suppression scaled with adaptive pressure, and decayed with a time course consistent with short-term synaptic plasticity (*10*). The change in activity could equally arise from deviation from the predicted stimulus pattern (*19*, *22*, *24*, *41*). Indeed, facilitation increased with the sound-intensity contrast between the adaptor and probe, whereas suppression was strongest at low contrast, as expected of prediction error that amplifies responses to unexpected sounds and suppresses expected ones (*15*, *41*). However, when we altered the global statistics of the sequence while keeping the local stimulus history the same, neither facilitation, suppression, nor stimulus-specific adaptation changed. Unexpected sound omissions also did not evoke spiking expected from comparison between prediction and the sensory input (*27*, *28*). Adaptation alone was sufficient to explain facilitation and suppression properties we report by scaling the response according to a divisive normalization model. In that model, adaptation depresses the sensory drive and normalization pool at different rates (*2*), producing a close-to-full range of modulatory effects we observed, and their reduction in background noise. Together, these results show that deviance signals in auditory cortical spiking can arise from canonical adaptive normalization without an explicit predictive model or input.

### A single adaptive mechanism produces both suppression and facilitation

Adaptation extends beyond suppression, and preceding stimuli can also facilitate responses in both auditory (*11*, *12*) and visual cortex (*13*). In the auditory cortex, adaptive suppression can be explained by adaptor-driven synaptic depression (*10*). Facilitation follows a similar time course but has been reported mainly when the adaptor differs spectrally from the probe sound (*11*). We show that facilitation also arises for spectrally identical sounds that differ only in intensity.

The facilitation and suppression we report carry signatures of adaptation. Both effects diminished when we reduced the adaptors from two to one, lowering adaptive pressure (Fig. 2C). They also diminished when the interval after the adaptor lengthened to 1 s, consistent with recovery from short-term synaptic depression (Fig. 2F) (*32*). Positive and negative probe contrasts differentially modulated facilitation in monotonic and non-monotonic cells (Fig. 3E), which can be explained by different adaptation to stimulus-intensity statistics in these two cell classes (*8*). Lastly, facilitation and suppression did not divide cells into anatomically segregated types (Fig. S1D–E); instead, the effects formed a continuous spectrum explained by adaptive normalization (Fig. 5).

### Adaptive normalization pool accounts for the contextual effects

Divisive normalization, first proposed to describe contrast-dependent gain in visual cortex (*42*), is considered a canonical cortical computation (*36*). When its sensory drive and normalization pool depress at different rates, it can produce a near-full spectrum of adaptor effects, from suppression to facilitation (Fig. 5). Facilitation arises when the pool depresses more than the drive, and balanced depression of both produces the weak suppression common in our data (Fig. 5).

The model treats the two depression rates as free parameters, but their interaction likely depends on sound intensity and probe contrast. One explanation for why facilitating and suppressing profiles scale differently with probe contrast (Fig. 3, S2) is that depression of the sensory drive is intensity-specific, while that of the pool is broad. At large probe contrast, the probe reactivates the broadly adapted pool and the weakly adapted sensory drive, so the response increases. At low probe contrast, the probe engages the depressed sensory drive. Suppression is therefore strongest at low contrast.

Facilitation and suppression relative to the rate-level baseline can thus arise from the same principles as SSA. SSA has been modeled as a depression of standard-specific channels that spares the deviant (*43*, *44*). An adaptive normalization model extends this principle to all inputs that participate in adaptive normalization, producing a continuum of effects ranging from suppression to facilitation.

Adapting inhibitory inputs to the auditory cortex can implement divisive normalization. The divisive effect is thought to arise from shunting inhibition or from scaling of balanced inhibitory and excitatory synaptic inputs (*36*, *45*). Inhibitory neurons shape sound-intensity tuning in auditory neurons (*46*, *47*), placing them in a position to provide divisive influence during contextual modulation (*48–50*). Their activity also changes with sound repetition (*51*, *52*), which could implement depression of the normalization pool in the model.

Background noise reduced both facilitation and suppression, which we modeled as reduced adaptor efficacy in depressing the drive and the pool (Fig. 6). The lower efficacy is consistent with the noise reducing sound-evoked responses (Fig. S6B) (*38–40*). Because adaptation is activity-dependent, the lower sound-evoked response in noise implies that the adaptor depressed the sensory drive and normalization pool less. Ongoing noise-driven activity would additionally depress both pathways, further reducing what the adaptor could add. Auditory cortex partly compensates for background noise by adapting to its statistics, yielding noise-invariant represen- tations of sounds (*39*, *53*). Our results suggest that this invariance does not extend to modulations by the temporal context.

Our probe differed from the adaptor in intensity, but the model is not specific to sound-intensity deviants. It models change in the response gain depending on how strongly the sensory drive and normalization pool pathways are depressed, not on the stimulus feature.

### Contextual deviance signals are decoupled from prediction error magnitude

Predictive models learn stimulus statistics to minimize the error between predicted and actual input. Our stereotyped and variable blocks differ in pattern and global statistics and should therefore lead to different predictive models. To exemplify how these differences impact prediction error, we considered normative models that can emerge by tracking the global pattern of the sequence, sound probability, or sound position within the chunk. Under each model, the prediction error differs between the two sequences in magnitude or direction, for the deviant or the standard (Table S1). Our data do not support these predictions. The statistical model showed that the deviant response did not change by more than 14% (95% HDI; Fig. 4C–D), and did not detectably change responses to the standard (Fig. 4D). This is consistent with reports that global deviants do not or only weakly modulate responses in primary sensory areas (*29*, *30*). In the auditory study (*29*), local frequency deviants elicited larger responses when the block made them rare rather than common, and the difference persisted across silent gaps between sequences that were longer than in our study. However, deviant density was not matched between blocks, so accumulating adaptation to the sound could have contributed to the finding (*29*).

We observed that responses to the same sound intensity were stronger when it was deviant than when it was standard (Fig. S4) (*5*; but see *34*). When the response to a frequency deviant exceeds that to the same stimulus in a control sequence, it has been taken as evidence of a prediction-specific input to the deviant, termed ‘true deviance’ (*22*, *24*, *31*, *54*). Such control removes suppression by repetition but imposes suppressive contexts of its own: a cascade sequence is predictable, which on a predictive account suppresses the response, and a many- standards sequence adapts responses through the intervening sounds. Here, we modified the many-standards sequence by lengthening the interstimulus interval beyond the range of forward suppression (*10*), which limited the adaptive effects, and resulted in a stricter baseline. By that criterion, facilitated profiles would be described as ‘true deviance’; nevertheless, their responses did not scale with global context affecting the prediction (Fig. 4).

Predictive coding frameworks propose that a distinct class of error neurons might signal prediction errors (*27*, *41*). Facilitated cells are a candidate for such a population because their responses increased with deviation from the preceding sound (Fig. 3). The direction of this scaling depended on the sign of the probe contrast, consistent with different cells signaling positive and negative prediction error (*27*). However, unlike prediction-error neurons, the facilitated cells did not modulate their responses across stimulus blocks to follow global sequence statistics (Fig. 4).

Hypothesized error neurons should respond when an expected sound is omitted, because the prediction is unmatched by sensory input (*27*, *28*). We found no omission responses above baseline firing rate. Omission activity also did not exceed post-stimulus rebound activity when no further sound was expected (Fig. S5). Reports of omission responses in sensory cortex differ from ours in two respects: (1) their designs did not separate omission from post-stimulus rebound; (2) the responses were not precisely time-locked to the omitted sound or appeared as slow ramping signals (*55–58*) and can be explained by intrinsic timing rather than prediction signals (*59*). Our data argue against error neurons in the primary auditory cortex that signal time-locked omissions.

We conclude that deviance signals in primary auditory cortex during passive listening are most parsimoniously explained by adaptation. If prediction contributes to the deviance signals, it depends solely on recent stimulus history, so that its effects converge with those of adaptation. Adaptive normalization is itself not inconsistent with predictive coding. It can be part of an inference over competing sensory causes, suppressing inputs that are already explained and amplifying deviant ones, thereby reducing redundancy (*60–62*). Shunting inhibition contributing to divisive normalization (*36*) can also gate prediction error in ways that would not necessarily affect firing rates (*63*). Lastly, deviance signals that depend on prediction error might emerge conditionally, for example during movement (*64*).

### Functional significance of contextual adaptation

Auditory responses adapt to the distribution of sound intensity (*4*, *8*, *65*), shifting rate-level profiles so that their dynamic range covers the most common intensities, improving their encoding (*4*, *8*) and discrimination (*66*). We find that adaptation also produces contextual modulation that amplifies deviance. Both facilitated and suppressed responses increased with the absolute intensity contrast between the probe and the preceding sound. The population therefore carries a graded deviance signal relative to the adapted state, and facilitation extends the range of responses to the deviant.

Because contextual modulation depends on how effectively the preceding sound depresses sensory drive and the normalization pool, it is vulnerable to background noise, and we predict that contextual modulation is similarly affected in hearing loss. This is because even though cortical responses can partially compensate for peripheral damage, their spike timing and synchrony do not recover (*67*), and these factors shape short-term synaptic depression. Consistent with this, adaptation to sound-intensity statistics is impaired in the auditory midbrain after noise-induced hearing loss (*68*). In addition, age-related hearing loss is accompanied by reduced inhibition across the central auditory pathway (*69*), which, through inhibitory activity in the normalization pool, would affect adaptation strength. Hearing aids may compound this deficit. They transform the auditory stream by applying dynamic range compression that reduces intensity differences and, as a result, reduces facilitation of responses to contrasting sounds.

Together, our results show that adaptation alone can account for contextual effects in auditory cortex spanning from suppression to facilitation. These include responses above those to the same sound in regular control, which have been taken as evidence of prediction error (*22*, *24*, *25*). Divisive normalization with the sensory drive and normalization pool depressing at different rates captured the continuum of contextual modulation. Facilitation scaled with the absolute contrast between the deviant and the preceding sound, but this was the only property it shared with prediction errors. Because adaptive normalization depends on the relative depression of its inputs, it is fragile. Background noise reduces contextual modulation, and we predict altered modulation in hearing loss. Preserving the intensity contrast between successive sounds may therefore be important for cortical salience of deviance.

## METHODS

### Animals

Seven adult C57BL/6J wild-type mice were used for the study (3 female and 4 male). Mice were obtained at 6–7 weeks of age from Charles River UK Ltd, and the recordings were performed at 10–11 weeks of age. Mice were housed with 2–4 cage-mates until the time of craniotomy for acute recordings, after which they were separated into individual cages. Mice were housed under a 12:12 h reverse light cycle, and experiments were performed during the dark phase. Mice had free access to food and water. All experiments were performed in accordance with the Animals (Scientific Procedures) Act 1986 (United Kingdom) and Home Office (United Kingdom) approved project and personal licenses held by the authors.

## Method details

### Surgeries

Mice underwent two surgeries: the first one to implant a stainless-steel headplate attached to the skull to enable head-fixation, and the other to perform craniotomy and prepare for acute electrophysiological recordings.

Surgeries were carried out following minimal standards for aseptic surgery on 8–9 week-old mice. Meloxicam (2 mg/kg intraperitoneal) was administered as an analgesic prior to surgery initiation. Mice were anesthetized with isoflurane (5% induction, 1–2% maintenance) mixed with oxygen as carrier gas (flow rate 1.0-2.0 L/min) and placed in a stereotaxic frame. After the skin incision, the skull was exposed, and Bregma and Lambda were aligned horizontally. A custom-made stainless-steel headplate was attached to the cranium using translucent dental cement (Super-Bond C&B, Sun Medical), and sites for later craniotomy above the auditory cortex were marked bilaterally at -2.9 AP, 4.9 ML (in mm from Bregma). A ground screw was implanted over the cerebellum. To secure the screw and protect the skull, the screw and skull were covered with a thin layer of dental cement. After the surgery, the mice were monitored daily for 3 days and administered oral analgesia (Metacam, Boehringer Ingelheim, 1 mg/kg). Mice recovered for 7 days after surgery before further experimental procedures.

Approximately 2 weeks following the first surgery, a bilateral craniotomy of approximately 1.5 mm diameter was performed to allow later acute electrophysiological recordings. The exposed dura mater was covered with Dura-Gel (Cambridge NeuroTech), and the site was sealed with a silicone elastomer (Kwik-Cast, WPI). Mice were allowed to recover for at least 16◻h before recordings began.

### Sound stimuli

Acoustic stimuli were presented to head-fixed mice inside the acoustic chamber. Stimuli consisted of a harmonic chord comprising 21 logarithmically spaced tones of equal amplitude spanning 2-32 kHz. Each chord lasted 80 ms, including 5-ms linear onset and offset ramps. The chords were organized into chunks of 1–5 sounds, presented with a fixed inter-stimulus interval (ISI) of 170 ms. Each chunk was followed by a randomized silent inter-sequence interval lasting 2–3 s.

Mice were habituated to head-fixation over 3 days, with the head-fixation period gradually increasing. Starting on the final habituation day and continuing for 3 consecutive days, mice were exposed to sound stimuli.

### Experimental paradigms during recording

During electrophysiological recordings, the following blocks of stimuli were tested:

1. Tones sequence. The recordings started and finished with blocks of pure-tone stimulus presentations. These were played in pseudorandom order across frequencies spanning 2–32 kHz or 2–48 kHz, spaced logarithmically, with a density of 5 frequencies per octave. Each tone lasted 100 ms and was presented with a 400 ms ISI at 55 dB or 75 dB SPL, 10 times per level. The tones were played with 5-ms-long linear onset and offset ramps.
2. Harmonic chords rate-level sequence. The chords were played in pseudorandom order across 30–85 dB SPL in 5 dB increments. Each chord lasted 80 ms and was presented with a 170 ms ISI, 10 times per level. The responses were used to assess cell monotonicity.
3. Interrupted oddball sequence (variable block). The stimulus sequence consisted of 3 chords presented with a 170 ms ISI. The first two chords were played at the same sound intensity (adaptor) and the third varied pseudorandomly across 7 levels (probe, 45–75 dB SPL in 5 dB increments). Each sound intensity occurred as the probe 15 times, resulting in a 4.8% probability of any given probe occurring across the block (calculated as 1/7 of the third-position chords). The adaptor intensity occupied 71.4% of the total density (the first two chord positions plus its occurrence at the third position). In each block, a single sound intensity was designated as the adaptor. The stimulus blocks described below systematically manipulated distinct features of the interrupted oddball sequence.
4. Harmonic chords rate-level baseline. Probe chords from interrupted oddball sequences were presented with a randomized ISI of 2–3 s without adaptors.
5. Stereotyped sequence. To evaluate the impact of standard density, a block of sequences with 95% standard density was presented. In 80% of chunks, these consisted of four sequential chords at the same sound intensity. In the remaining 20% (n=15), the third sound in the sequence was set to a deviant sound intensity, yielding an overall deviant density of 5% and a standard density of 95%.
6. Short interrupted oddball sequences. Sequences consisted of two chords, with the first serving as the adaptor and the probe varying across the 7 sound intensities, each tested 15 times.
7. Variable ISI oddball sequences. To test the effect of stimulus timing, the sequence was modified so that the ISI of the probe sound was pseudorandomly varied to 150 ms, 350 ms, or 1 s, each tested 15 times.
8. Noise-masked sequences. Background white noise was played at two different levels during the interrupted oddball sequence: 38 or 43 dB SPL (low noise) and 53 or 58 dB SPL (high noise). The noise was linearly ramped from 0 to the peak intensity over a 0.35 s window beginning at the onset of the first adaptor in the sequence. The peak noise level was reached at the offset of the second adaptor and was sustained for 0.25 s following the probe offset.
9. Omission sequences. To test neuronal activity during omission of expected sound, the stereotyped sequence was modified to have 5 chords per chunk. It had no sound deviants but omitted the third sound in 20% of chunks (n=15). The test was performed in a separate group of mice, distinct from those exposed to interrupted oddball sequences with 71% standard density.

### Sound delivery

Sounds were delivered using MATLAB (MathWorks) and a digital signal processor (RZ6, Tucker-Davis Technologies). They were presented in an acoustic chamber (IAC Acoustics) via an open-field magnetic speaker (MF1, Tucker-Davis Technologies) directed at the left or right ear and placed 10 cm from the ear opening. Before a batch of experimental animals was introduced to the acoustic chamber, the acoustic stimuli were calibrated using a microphone (480C02, PCB Piezotronics) positioned at the approximate location of the animal’s ear opening during the recording sessions. The calibration was performed using pure tones in the 2–48 kHz range to ensure the sound system’s frequency response was flat to within ±3 dB.

### Electrophysiology recording

Acute electrophysiological recordings were performed in awake, head-fixed mice using Neu- ropixels 1.0 or 2.0 probes (IMEC) from the hemisphere contralateral to the ear closer to the speaker. The probes were coated with DiO, DiI, or DiD cell-labeling solution (Thermo Fisher; V22889) before each recording to allow histological identification of probe tracks. Each record- ing from the same hemisphere used a different cell-labeling solution, with up to three recordings in the hemisphere.

During the recording session, mice were placed in a custom cylindrical tube inside the sound chamber and head-fixed to a custom mounting clamp with two screws. Silicone elastomer (Kwik- Cast, WPI) protecting the craniotomy was removed. The probe’s ground wire was connected to the implanted ground screw. The probe was positioned to target auditory cortex and was lowered perpendicularly below the brain surface using a micromanipulator (uMp-4, Sensapex) to a depth of 1.4–2.0 mm with a speed of 3–5 µm/s. In a subset of recording sessions, a second Neuropixels probe was inserted in the medial geniculate body of the thalamus (thalamic data not reported here). To ensure recording stability, the probe was allowed to settle in the tissue for 25–30 min after insertion prior to testing. Left and right hemispheres from the same mice were recorded in 1–3 recording sessions each, depending on the ability of the probe to penetrate the brain via the same craniotomy on a different recording day.

Electrophysiological data were acquired at a 30 kHz sampling rate; when using Neuropixels 1.0, an additional data stream at 2.5 kHz for the LFP band was recorded. Probes were connected via the PXIe card inside a National Instruments chassis, and Open Ephys GUI was used for acquisition. The probes were configured to use ground as the reference.

For Neuropixels 2.0 configurations, where the total number of physical recording sites exceeds 384 channels capable of simultaneous recording, the active channel map was customized online to maximize the count of channels within the auditory areas. To achieve this, recording sites across the shanks spanning the entire length of the probe inserted in the brain were surveyed for pure tone responses (2 kHz to 48 kHz) and to the harmonic chords across sound pressure levels. Evoked spiking activity and LFP signals were used to identify continuous probe segments that responded to the stimulus and were included in the final channel map.

In addition to electrophysiology data, video of the animal and stimulus parameters and timing were recorded. The video camera (Firefly FFY-U3-04S2M, Flir) was directed at the animal’s face and recorded pupil changes and face movements at a frequency of 30 frames / s. Low- intensity infrared light illuminated the chamber and the mouse face. The stimulus data streams included stimulus onset, offset, and identity and were recorded through Open Ephys Acquisition Board. The stimulus data streams and electrophysiological signals were synchronized within Open Ephys GUI. The synchronization pulses were provided by Arduino Uno sending a periodic TTL signal (period 1 s, random duration of on period 200–300 ms) to PXIe card and to the video camera, which registered its status alongside each video frame.

### Histology

Following the electrophysiology recordings, animals were terminally anesthetized by intraperi- toneal injection of pentobarbital and then transcardially perfused with phosphate-buffered saline (PBS) followed by 4% paraformaldehyde (PFA). Brains were removed and post-fixed for 24–48 hours, then rinsed and subsequently cryoprotected overnight in 30% (w/v) sucrose dissolved in PBS. Coronal brain sections were cut using a microtome (Leica) with 80–100 μm thickness. After rinsing in PBS, the sections were mounted in Fluoroshield with DAPI (Sigma). The sections were inspected to locate Neuropixels probe tracks.

### Electrophysiological data pre-processing and spike sorting

Action potential data streams recorded from Neuropixels probes were pre-processed using the SpikeInterface Python package. The streams were high-pass filtered (300 Hz) and corrected for hardware-sampling offsets (SpikeInterface *phase_shift* function). Bad channels were detected and excluded based on outlier standard deviations, low similarity to neighboring channels, or excessive high-frequency power (default parameters of SpikeInterface *detect_bad_channels* function). The processed data streams from a single session were then merged immediately prior to spike sorting.

Automated spike sorting was performed on the merged data using Kilosort 2.5. Prior to spike sorting, the software performed pre-processing steps, such as drift correction and signal whitening. The spike-sorting results were processed using Bombcell software to calculate single-unit and template quality metrics. Manual curation was subsequently performed in Phy on the extracted clusters: oversplit clusters of putative single units were merged, and clusters were classified as good units, multi-unit activity or electrical noise. The classification was based on visual inspection of waveforms, cluster stability, and the quality metrics provided by Bombcell and Kilosort.

### Cell type classification

Curated single units were imported into CellExplorer for final validation and physiological classification. Based on extracellular waveform shape (trough-to-peak duration and asymmetry) and the unit’s spike time autocorrelogram, CellExplorer classified units as pyramidal cells, narrow interneurons, or wide interneurons.

### Electrophysiological data analysis

#### Unit inclusion criteria

Single units included in the analysis had to meet criteria for firing rate, stability, and sensory responsiveness. Because individual units could drift or experience signal loss during the acute recordings, inclusion criteria were applied independently for each experimental epoch. An experimental epoch was designed to contain all stimulus conditions across which direct firing rate comparisons were made (block 1: data for Fig. 1, 2A–C, 3, 4, 5; block 2: data for Fig. 2D–F; block 3: data for Fig. 6).

Within a given experimental epoch, the included single units had to satisfy the following:

1. Firing rate: Units had to maintain an overall mean firing rate ≥ 0.5 Hz.
2. Temporal stability: Units had to maintain a stable firing rate as assessed by Gini coefficient (FR_gini_ ≤ 0.6). Gini was calculated across firing rates estimated in consecutive 2-minute intervals throughout the block.
3. Chord-responsiveness: A unit was classified as chord-responsive when it had a significant stimulus response for at least one sound level. A significant stimulus response was defined by increased firing rate during the 80 ms chord presentation relative to an 80 ms pre-stimulus baseline (p < 0.05, Wilcoxon signed-rank test), while also exhibiting spikes in at least 33% of the trials at that specific sound level. P-values for the same cell across sound intensities were adjusted using the Benjamini–Hochberg false discovery rate correction.

#### Identification of the primary auditory area

Recording sites within primary auditory cortical areas (primary auditory cortex, A1; anterior auditory field, AAF) were identified by electrophysiological criteria and the probe tracks were histologically verified. To delineate the spatial extent of the cortex along the probe, mean tone-evoked local field potentials (LFPs), multi-unit activity (MUA) peri-stimulus time histograms (PSTHs), and the LFP-derived current source density (CSD) profile were calculated and assessed. They were used in corroboration to identify the extent of the cortical area across the probe depth, as well as the depth of the boundary between granular and infragranular layers.

To distinguish shank insertions within primary auditory areas (AAF and A1) from those in secondary areas, tone-evoked responses were assessed with a method adapted from previous studies (*70*, *71*). Given the perpendicular insertion of the probe relative to the brain surface, all recording sites spanning a single shank at the depths within the cortex were assumed to be located in the same cortical area. Multi-unit spiking activity recorded on each channel (extracted via Kilosort) was evaluated for tone-evoked responses. The responses to tones of different frequencies and attenuations were pooled and compared to the preceding baseline (p < 0.05, Wilcoxon signed-rank test).

For a shank location to be classified as within the primary auditory cortex, its units had to satisfy two criteria. First, at least 25% of the cortical units had to exhibit a significant tone response. Second, the tone-responsive units were required to exhibit response latency ≤ 20 ms. Response latency was calculated as the first 2-ms bin following stimulus onset in which the tone-evoked MUA crossed a threshold of 2 standard deviations above the baseline mean (*72*). The 25th percentile of the response latencies across the shank’s units was taken as its statistic.

### Laminar estimation

Laminar position along each shank was estimated from pure tone responses. Two complementary measures were combined: the current source density (CSD) profile of the LFP, and the depth profiles of multi-unit activity (MUA). The estimation was performed separately for each shank and recording session. Its output was the depth of the boundary between the granular and infragranular layers, which served as the reference 0 µm.

The depth of the granular–infragranular boundary was assigned by corroborating three markers. The boundary had to lie: (1) below the earliest current sink in the CSD, taken to mark the granular layer, (2) at or below the shortest-latency MUA, (3) above the depth of peak MUA, taken to mark layer 5. Each unit’s depth was defined as the position of the channel with the largest waveform amplitude. Positive depths indicate positions below the granular-infragranular boundary.

### Spiking activity metrics

To quantify changes in neural responses, the following were calculated on sound-evoked firing rates: Modulation index (MI): Measures the change in response to stimulus condition A relative to stimulus condition B.

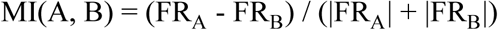

Facilitation index of a profile was calculated on spontaneous rate-corrected firing rates as modulation index of the mean adapted relative to the mean baseline response:

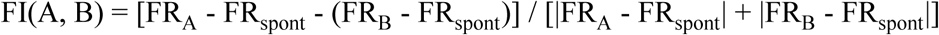

Common SSA index (CSI): Measures the overall strength of stimulus-specific adaptation across symmetric choices of deviant and standard stimuli (*33*). For a low sound level (L_low_, 50–55 dB SPL) and a high sound level (L_high_, 75 dB SPL), the index combines their respective standard and deviant responses as follows:

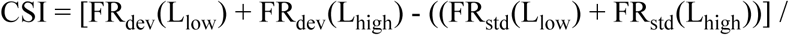

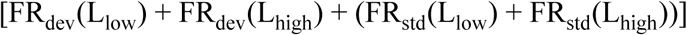

Monotonicity index: quantifies rate-level profile monotonicity, calculated on the profile’s firing rate above baseline as the firing in response to the highest-intensity sound over the profile’s maximum firing (*8*). Profiles with monotonicity index > 0.5 were classed as monotonic, and those ≤ 0.5 as non-monotonic. The index was quantified on the cell’s rate-level baseline. Cells that were not classified as chord-responsive during the rate-level baseline had undefined monotonicity and were excluded from the analysis in Fig. 3.

Dynamic range of facilitation/suppression: the longest continuous range in the rate-level profile over which the responses increased (facilitation) or decreased (suppression).

### Analysis

When presenting PSTH responses across the cell population (Fig. 1H and S5B), firing rates were binned with 2.5 ms bins, smoothed with a Gaussian kernel (σ = 2.5 ms), and normalized to the cell’s maximum smoothed firing rate across all probe stimuli.

For z-scoring neuronal activity, spontaneous activity before the chunk onset was used as a baseline.

#### Adaptive normalization model

The computational model is derived from the divisive normalization model (*36*). The rate-level baseline response can be described as

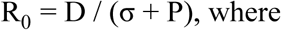

D is the sensory drive, P is the strength of the normalization pool, and σ is the semi-saturation constant, which also prevents the denominator from approaching zero.

The adaptor depressed the drive and the normalization pool by fractions α_D_ and α_P_, respectively, resulting in the adapted response:

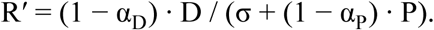

The response gain is therefore independent of D:

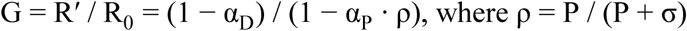

G was computed for α_D_ and α_P_ combinations ranging from 0 to 1 in steps of 0.01, with σ = 0.1 and P = 1.0. Because σ and P enter the gain through ρ, changing their values changes the slope of the G = 1 line, but they result in qualitatively the same results.

The model can produce full suppression for α_D_ = 1, so G_min_ = 0, FI_min_ = -1.

The maximum facilitation is bounded by ρ. It is maximized for α_D_ = 0 and α_P_ = 1: G_max_ = 1/(1 − ρ)

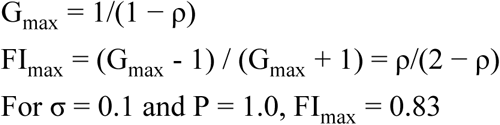

To simulate background noise, both depression rates were scaled by the adaptor’s efficacy k:

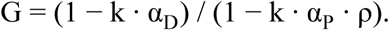

Gain was computed for the following pairs of α_D_ and α_P:_ α_D_ = [0.3, 0.4, 0.5, 0.6, 0.7], α_P_ = [0.8,

0.7, 0.55, 0.4, 0.3]. The simulation considered two scenarios: increasing σ from 0 to 1 with k = 1, and decreasing k from 1 to 0 with σ = 0.1.

## Statistical analysis

Unless stated otherwise, data are reported as mean ± SEM.

### Identification of facilitated and suppressed profiles and responses

To identify instances in which the rate-level profile changed significantly following the adaptor relative to the baseline, the effect of each adaptor intensity was assessed separately with a permutation test. To create a null distribution of expected changes by chance, the label of each trial (baseline vs adapted) was shuffled. For each shuffle, the change from the rate-level profile was calculated as a sum-squared error. A change in the adapted rate-level profile from the baseline profile was classified as significant when its magnitude exceeded that in 95% of the shuffled profiles (1000 shuffles). For each adapted profile, a facilitation index was calculated. Those profiles that changed significantly and had a positive facilitation index were classified as facilitated; those that changed significantly and had a negative facilitation index were classified as suppressed.

A response was classified as facilitated if it belonged to a facilitated profile and exceeded the rate-level baseline at that intensity. It was classified as suppressed if it belonged to a suppressed profile and fell below the rate-level baseline at that intensity. All other responses were classified as without significant change.

For comparison of adaptation effects across different conditions, the profiles were classified as facilitated or suppressed on a subset of data, and the held-out data were used for calculation of the compared metric. This approach ensures that selection criteria for the profiles do not bias the comparison results. For data in Fig. 2C, the profiles were classified as facilitated or suppressed on probe intensities of 50, 60, and 70 dB SPL and the remaining probe levels were used for calculation of response compared in Fig. 2C. For data in Fig. 4, the profiles were classified on probe intensities other than those of deviants and standards in the stereotyped block, and the responses of those profiles were compared for held-out probe intensities.

In some experimental manipulations of the sequence, data for a subset of probe intensities from the range of 45–75 dB SPL were collected. In these cases, only the intensities present in these manipulations were included in the data used for classification (data in Fig. 2F and Fig. 6).

### Mixed effects models

To account for dependencies between data samples, significance tests were performed on mixed- effects models (*lme4* and *lmerTest* packages for linear models, *glmmTMB* for generalized models in R). Random intercepts were included for cell IDs nested within recording sessions to account for their correlations. For models with multiple fixed effects, the interactions were tested, and the reported model includes only significant interactions. Linear mixed-effects models were tested with F-tests after Satterthwaite’s approximation for degrees of freedom. Poisson mixed models were tested with a Wald χ² test.

Post-hoc analysis on the linear models was performed for the pairwise effects and slopes per group using *emmeans* package in R and Kenward–Roger approximation for degrees of freedom. P-values were adjusted with Bonferroni correction for multiple comparisons.

### Bayesian hierarchical model

All models were fit in R using *brms* package. Their convergence was assessed using R-hat values and effective sample size. Spike-count models (Fig. 4, 6C) used a Poisson likelihood on spike counts summed across trials. Effects were estimated on the log scale and gain is reported as their exponential (gain of 1 indicating no scaling). Group-level effects were nested across session, cell, and adaptor intensity to account for the correlations between samples. Priors were weakly informative: Normal(0,1) for population-level effects on the log scale, half Student-t(3, 0, 1) for group-level standard deviation. Posterior estimates for the gain are reported as the median and the 95% highest density interval (HDI) — the narrowest interval containing 95% of the posterior probability mass.

Response gain in Fig. 4: The models included a fixed effect of block (variable vs stereotyped) that varied across session, cell, and adaptor. The model of the response to the first standard also included the number of standard repetitions in the previous chunk as a continuous predictor. Facilitated and suppressed profiles were modeled separately. The total deviation of a profile from its group-level gain was calculated as the sum of its posterior random-effect estimates across the session, cell, and adaptor level. To consider the profile’s gain to be higher than that estimated for the group, its 95% HDI gain calculated from the profile’s total deviation had to be entirely above 1.

Response gain in Fig. 6C: The spike counts were modeled as the sum of baseline and evoked firing to separate the adaptor’s effect on evoked rate from those due to the effect of noise on baseline firing.

CSI shift in Fig. S4: The difference in CSI between two sequence blocks was modeled with a Student-t likelihood and an intercept representing the mean shift. Each cell’s standard error, estimated by bootstrapping across trials, was supplied as measurement error.

## ACKNOWLEDGEMENTS

We thank Jennifer Linden and Sam Solomon for their valuable feedback on the manuscript.

## Funding

RNID Fellowship: FG24/9 (P.J.), Biotechnology and Biological Sciences Research Council (BBSRC) Research Grants: BB/Z51665X/1, BB/Y010345/1 (D.B.)

## Author Contributions

P.J. and D.B. designed the research; P.J. and A.B. performed the research; P.J. analyzed the data and wrote the paper with support from the other authors; all authors approved the final version of the manuscript.

## Data and code Availability

Aggregated spike data will be made publicly available upon publication. Because of the dataset size, the raw electrophysiological and spike-level data will be available upon request. Analysis code is available from the authors’ GitHub at https://github.com/bendor-lab/auc_adapt_norm_pool.

## Declaration AI-assistance

During the preparation of this work, the authors used AI tools for coding assistance (Claude), manuscript editing (Claude and Grammarly), and feedback (Claude). The authors take full responsibility for the final content of the manuscript.

## Declaration of interests

The authors declare that they have no competing interests.

## Notes

### Competing Interest Statement

The authors have declared no competing interest.

