## Supplemental Materials for "Adaptation can explain cortical responses to sound deviance without requiring prediction"

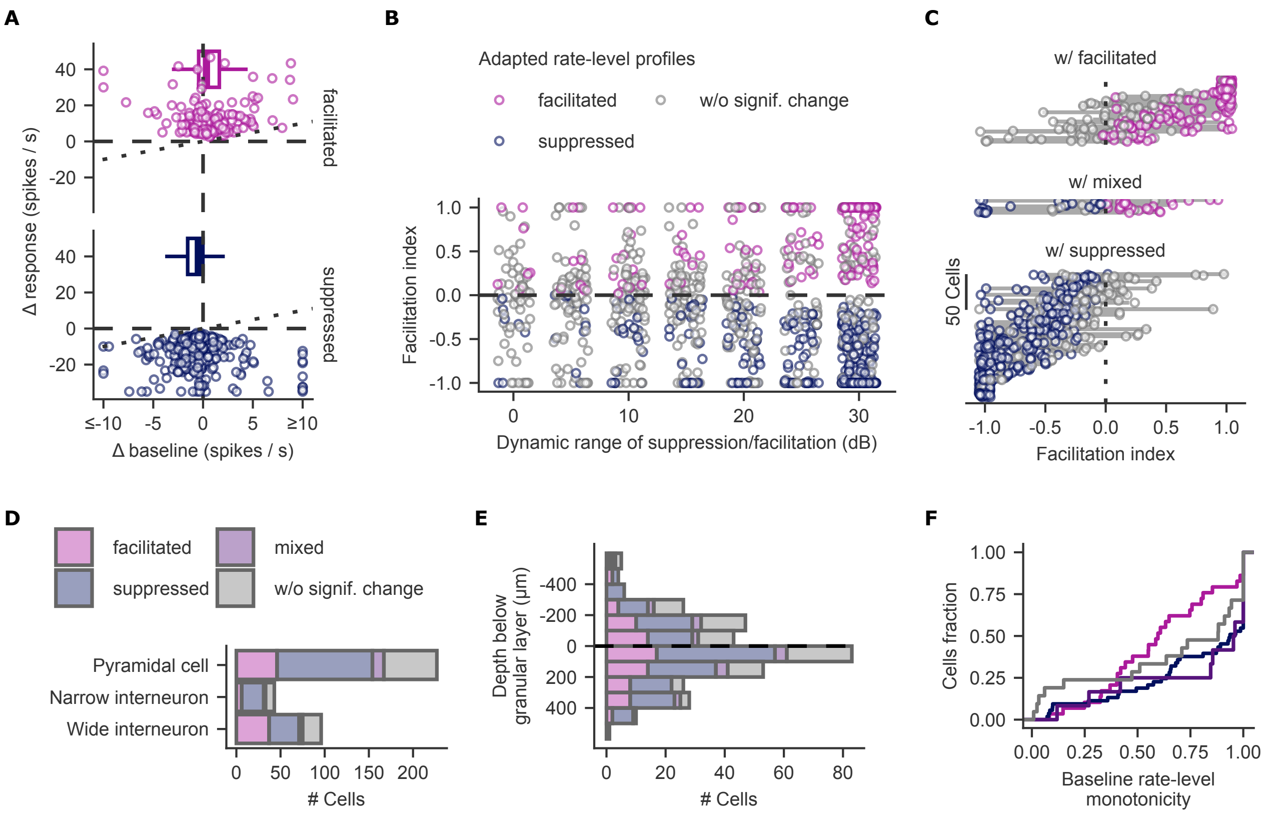


**Fig. S1: Effect of sound context on spiking across cell types, cortical depth, and monotonicity profiles. Related to Fig. 1.**

A. Change in mean probe response within a given profile plotted against the change in spontaneous activity for facilitated and suppressed profiles. We calculate the change in spontaneous activity (Δbaseline) as the firing rate during the 80 ms window preceding the probe onset minus that during the window preceding the first adaptor. Dotted lines mark equality of the change. Data for facilitated profiles above the line signify higher change in the evoked response than to the baseline; for suppressed profiles the change to the evoked responses is lower than that to the baseline. Boxplots summarize Δbaseline (Boxes show the median, IQR, and whiskers extend to 1.5 × IQR). A linear mixed-effects model showed that Δbaseline differed significantly between facilitated and suppressed profiles: $F_{(1, 392.7)} = 18.9$, $p < 10^{-4}$. Estimated Δbaseline for facilitated profiles was 1.0 spikes/s (95% CI [0.1, 1.9] spikes/s); for suppressed profiles it was -0.3 spikes/s (95% CI [-1.1, 0.4] spikes/s).

B. FI of a profile plotted as a function of its dynamic range of adaptation. Data for all facilitated, suppressed, and non-significantly changed profiles. Points are jittered horizontally for visibility.

C. FI computed across different adaptors tested on the same cell. Each row represents data for a single cell; horizontal lines connect its data. For visualization, cells are grouped by adaptation class: cells with a facilitated profile (n=89), cells with a suppressed profile (n=173), cells with both (mixed, n=19). Within each adaptation class, cells are sorted by median FI. In cells with a facilitated profile, 60 ± 5% of the remaining profiles also had a positive FI; in cells with a suppressed profile, 78 ± 3% of the remaining profiles had a negative FI.

D. Count of cells in each adaptation class by putative cell type.

E. Depth distribution of recorded cells in each adaptation class relative to the granular-infragranular boundary. Histograms are binned in 100 μm bins.

F. Cumulative fraction of cells in each adaptation class as a function of monotonicity index. The monotonicity index was calculated on the cell's rate-level baseline.

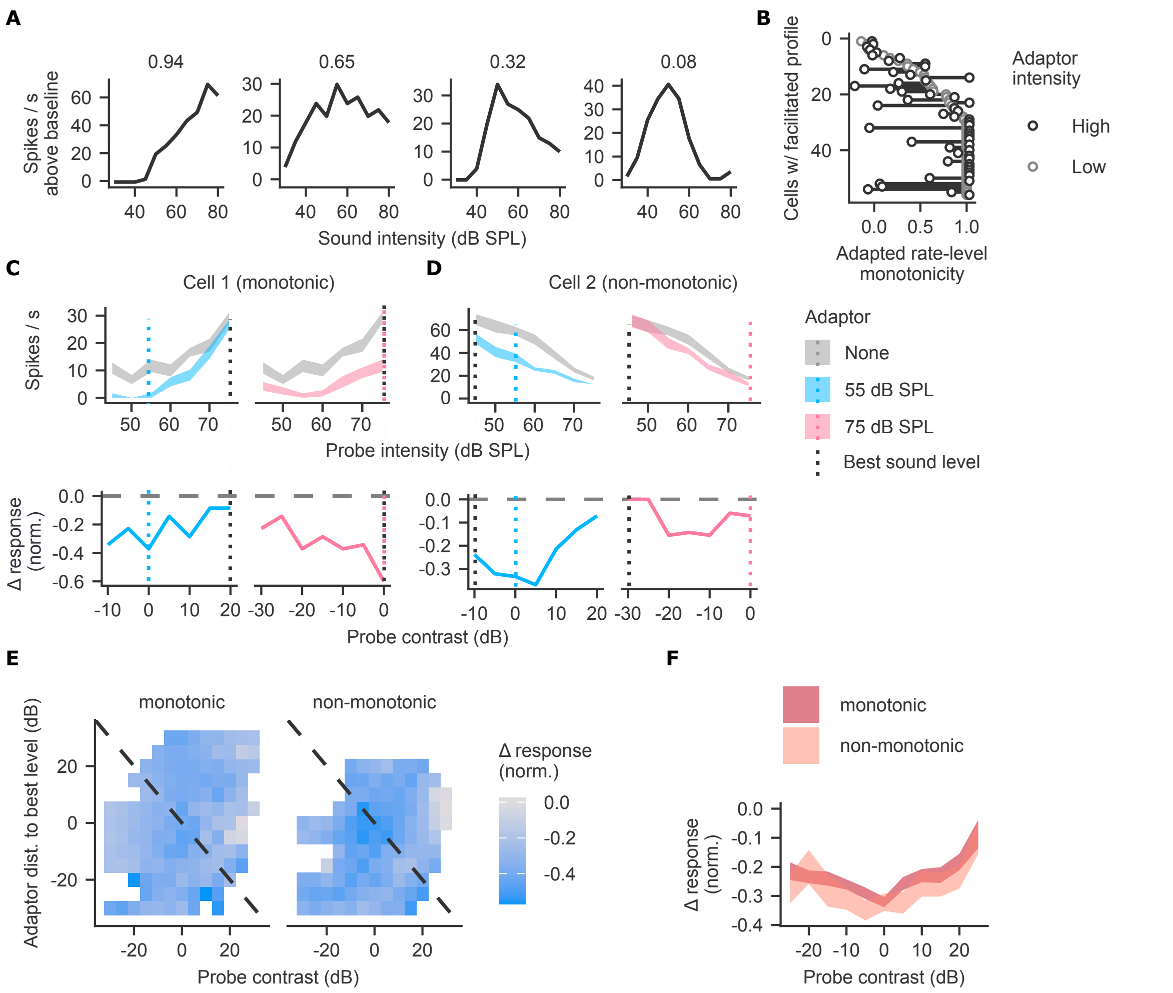


**Fig. S2: Classification of cells by sound-level monotonicity, and the contrast dependence of response suppression. Related to Fig. 3.**

A. Example rate-level profiles calculated on harmonic chords sequences for cells of different monotonicity. The cell's monotonicity index is shown above its rate-level profile.

B. Monotonicity index of cells with a facilitated profile compared between profiles that follow a low- and high-intensity adaptor. Data shown only for cells responsive in both conditions (n=56) and not shown for suppressed profiles due to their low spiking rates following adaptor making the monotonicity index unreliable. Monotonicity index change of facilitated profiles: 0.22 ± 0.04.

C–D. Example monotonic (C) and non-monotonic (D) cells with a suppressed profile plotted as in 3B–C.

E. Mean normalized response change as a function of probe contrast and of the adaptor's distance from the best level, for monotonic and non-monotonic cells plotted as in 3D.

F. Response change as a function of probe contrast plotted as in 3E.

A linear mixed-effects model of suppression magnitude (sign-inverted normalized response change, so positive values indicate stronger suppression) was fitted analogously to Fig. 3 (n = 2,588 responses from 375 profiles from 125 cells in 17 sessions; includes only chord-responsive cells during harmonic chords rate-level sequence). The effect of positive probe contrast × monotonicity: $F_{(1, 2562.3)} = 0.6$, p = 0.4; negative probe contrast × monotonicity: $F_{(1, 2577.4)} = 5.5$, p = 0.02. For monotonic cells, the scaling with positive contrast was −0.0060, 95% CI [−0.0076, −0.0043] per dB; with negative contrast, −0.0057, 95% CI [−0.0072, −0.0041] per dB. For non-monotonic cells, the scaling with positive contrast was −0.0072, 95% CI [−0.0097, -0.0047] per dB; with negative contrast, −0.0020, 95% CI [−0.0046, 0.0006] per dB. The effect of adaptor distance × monotonicity: $F_{(1, 1733.8)} = 15.0$, $p = 10^{-4}$. Slopes per dB were -0.0025 [-0.0039, -0.0012] for monotonic cells and 0.0023 [0.0002, 0.0044] for non-monotonic cells.

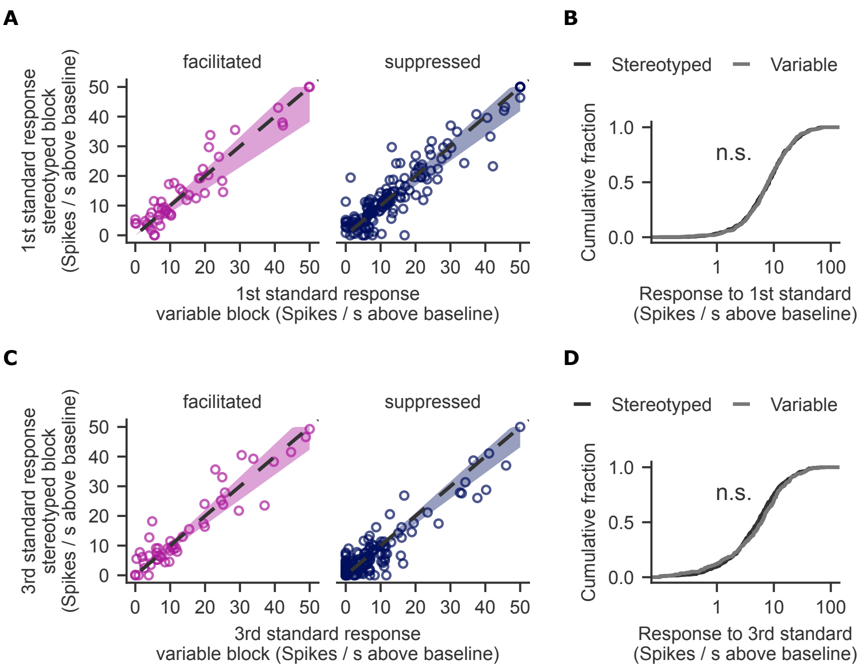


**Fig. S3: Responses to standard sounds do not scale with sequence statistics when local history is matched. Related to Fig. 4.**

A. First-standard responses in the stereotyped versus the variable block. The first standard followed 2–3 s of silence in both blocks. Data for first-standard responses significantly exceeding spontaneous activity in either block, facilitated responses: n = 67 from 46 cells; suppressed responses: n = 168 from 93 cells. The dashed line marks equality; the colored band shows the 95% HDI of the gain from a Bayesian hierarchical model that included the number of standards in the preceding chunk as a covariate. Gains are given in Fig. 4D.

B. Cumulative distributions of first-standard responses in the two blocks. Data for responses significantly exceeding spontaneous activity in either block (n = 460 responses per block from 228 cells). Kolmogorov–Smirnov test, p = 0.83.

C. Third-standard responses in the stereotyped versus the variable block for profiles whose responses are included in A. The dashed line marks equality; the colored band shows the 95% HDI of the gain from a Bayesian hierarchical model. Gains are given in Fig. 4D.

D. Cumulative distributions of third-standard responses in the two blocks for profiles whose responses are included in B. Kolmogorov–Smirnov test, p = 0.10.

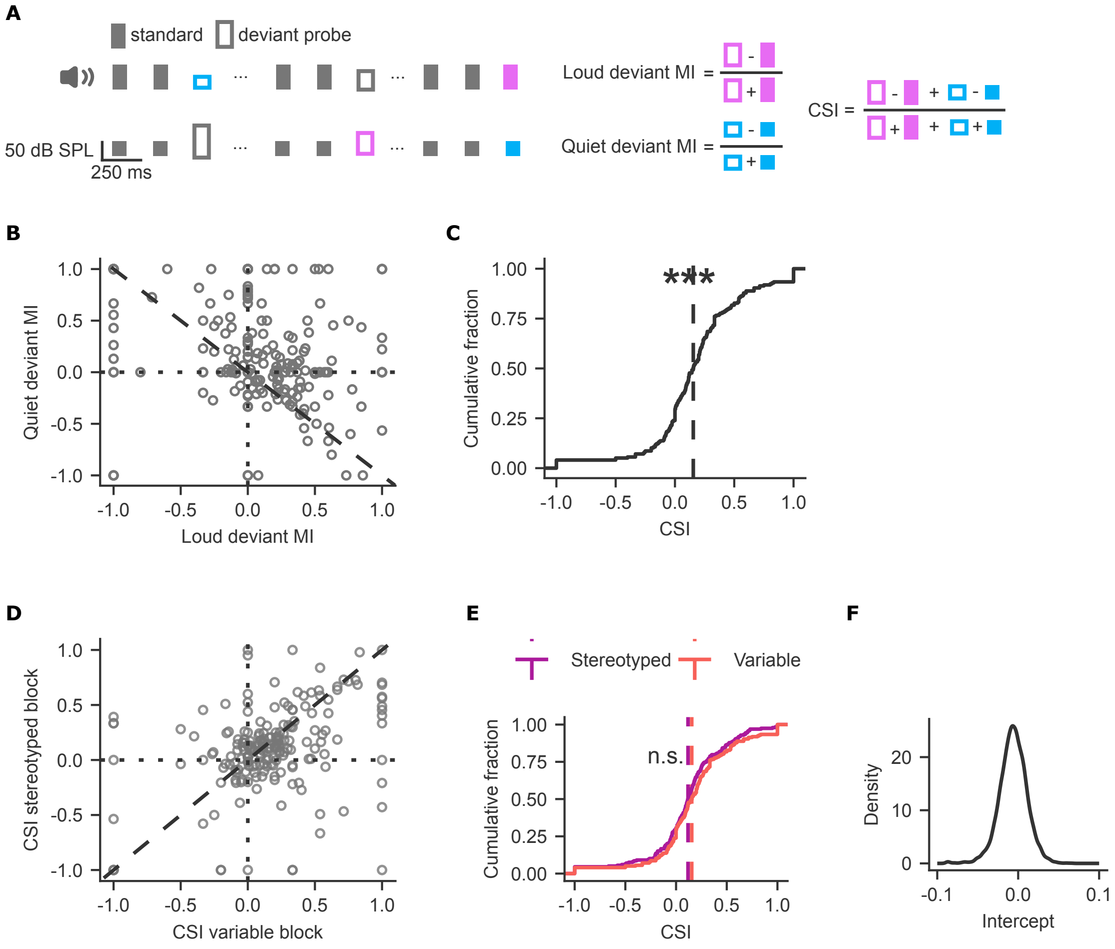


**Fig. S4: SSA does not scale with standard density. Related to Fig. 4.**

A. Calculation of the deviant modulation indices (MIs) and the common SSA index (CSI).

B. Quiet deviant MI plotted against loud deviant MI for all chord-responsive cells in the variable block (cells as in Fig. 4E). Dotted lines mark zero MI. The dashed line marks where the two MIs average to zero. Cells above that line have a positive mean MI.

C. Cumulative fraction of CSI for the cells in B. Vertical dashed lines indicate the median. Wilcoxon signed-rank test for median of 0: V = 14549, $p < 10^{-11}$.

D. CSI in stereotyped block plotted against the variable block for the same cells. Dotted lines mark zero, and the dashed line marks equal CSI.

E. Cumulative fraction of CSI for the same cells in the stereotyped and variable blocks. Vertical dashed lines indicate the respective population medians. Kolmogorov-Smirnov test, p = 0.27.

F. Posterior density for the shift in CSI between stereotyped and variable blocks. The shift was estimated as the intercept of a Bayesian hierarchical errors-in-variables model; median -0.01, 95% HDI [−0.04, 0.03].

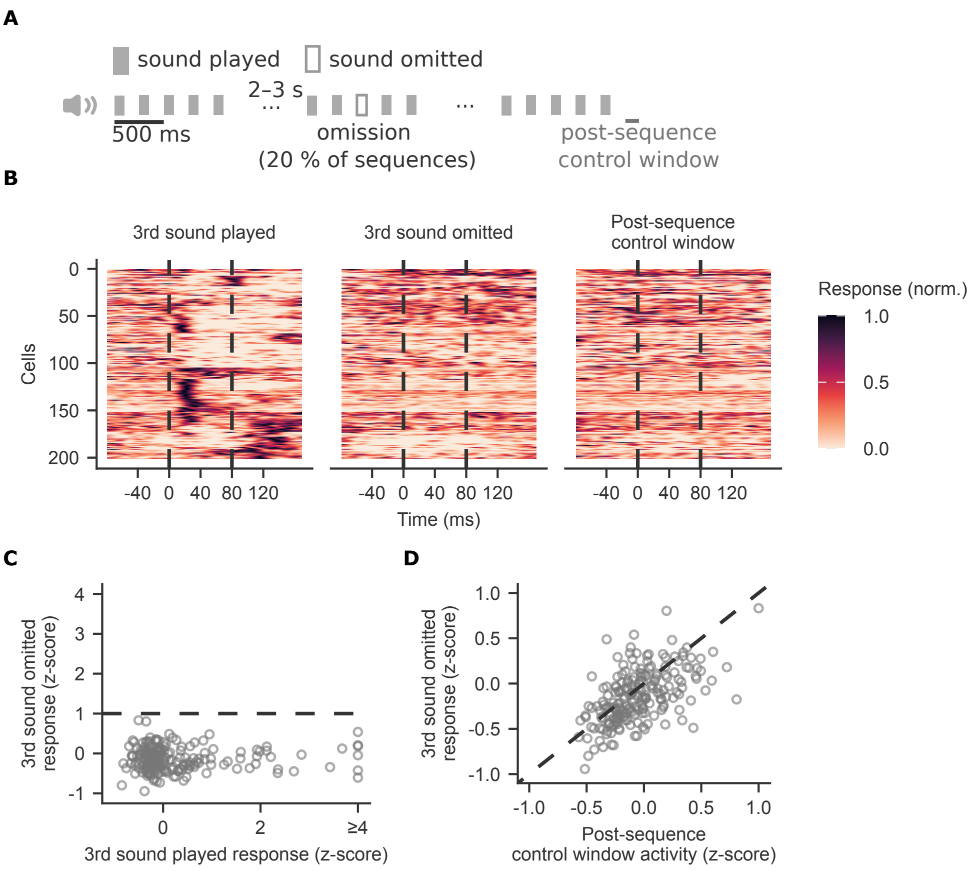


**Fig. S5: Unexpected sound omission does not evoke a spiking response. Related to Fig. 4.**
A. Schematic of the stimulus paradigm in which the 3rd sound is omitted in 20% of sequence chunks.
B. Normalized firing rate of each cell (one cell per row; n = 201 cells) aligned to the onset of the 3rd sound (left), the expected onset of the omitted 3rd sound (middle), and the onset of the post-sequence control window (right) as shown in A. We normalized firing rates to each cell's maximum across the three conditions. Cells are sorted by hierarchical clustering and displayed in the same order across panels. Dashed lines mark the onset and offset of the 80 ms sound (or its unexpected omission).
C. Response during the omission window plotted against the response to the played 3rd sound. Responses are z-scored relative to the spontaneous activity during the 80 ms window preceding chunk onset. The dashed line marks a z-score of 1.
D. Activity in the omission window plotted against activity in the post-sequence control window. The post-sequence control window shown in A is aligned to where a sixth sound would have occurred. The dashed line marks equality. The difference between omission and control window activity did not exceed a z-score of 0.31 in 95% of cells.

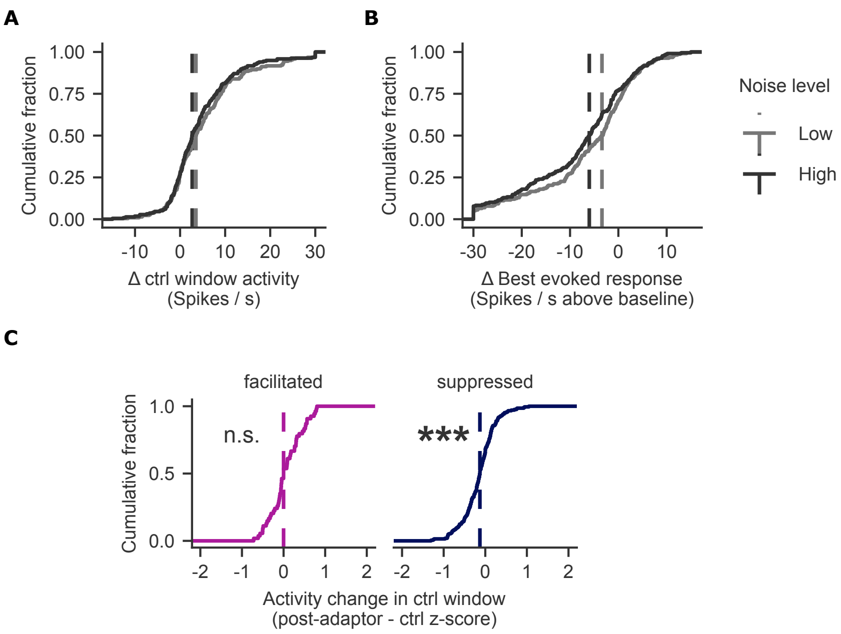


**Fig. S6: Effects of background noise on adapted responses. Related to Fig. 6.**

A. Cumulative fraction of the changes in spontaneous firing rate in low and high noise relative to no noise. The firing rate is calculated on the control window without an adaptor, as shown in Fig. 6A. Dashed lines mark the medians. Data for cells in low or high noise, as in Fig. 6C–D. Noise increased baseline activity at both noise levels (Wilcoxon signed-rank tests): low noise median change: 3.5 spikes / s, V = 19842, $p < 10^{-16}$; high noise median change: 2.7 spikes /s, V = 19348, $p < 10^{-15}$.

B. Cumulative fraction of the changes in best evoked response in low and high noise relative to no noise. The best evoked response is the maximum response across the four probe intensities presented in noise. The change is the response in noise minus the response without noise, without an adaptor. Dashed lines mark the medians. Same cells as in A. Noise reduced the best evoked response at both noise levels (Wilcoxon signed-rank tests): low noise median change: -3.4 spikes/s, V = 4876, $p < 10^{-13}$; high noise median change: -6.0 spikes/s, V = 3527, $p < 10^{-16}$.

C. Change in pre-probe activity after the adaptors. A linear mixed-effects model showed a significant difference between facilitated and suppressed profiles: $F_{(1, 86.7)} = 11.8$, $p < 10^{-3}$. Data are for facilitated (n = 18) and suppressed (n = 69) profiles, pooled across noise levels. Estimated marginal means were tested against zero: for facilitated profiles: 0.06 z-score (95% CI [−0.06, 0.18]); for suppressed profiles: −0.15 z-score (95% CI [−0.21, −0.09]).

| Predictive model | Model description | Quantity setting the prediction error | | Deviant | | | 1st standard | | | 3rd standard | | |
|---|---|---|---|---|---|---|---|---|---|---|---|---|
| | | Description | Formula | Var[a] | Ster[b] | Gain | Var[a] | Ster[b] | Gain | Var[a] | Ster[b] | Gain |
| Global rule | Learned how often the third sound differs from the first two in the block[c] | Information content of a deviant at 3rd position | $-\log_2$ P(change) | 0.22 bits | 2.32 bits | >1 | — | — | ~1[d] | 2.81 bits | 0.32 bits | <1 |
| Positional probability | Probability of each sound level at each position in the chunk | Information content of the sound level at its position | $-\log_2$ P(level \| position) | 2.81 bits | 2.32 bits | <1 | 0.0 bits | 0.0 bits | ~1 | 2.81 bits | 0.32 bits | <1 |
| Density | Probability of each sound level in the block, irrespective of position | Information content of the sound level | $-\log_2$ P(level) | 4.39 bits | 4.32 bits | ~1 | 0.49 bits | 0.07 bits | <1 | 0.49 bits | 0.07 bits | <1 |
| Mean | Expected (mean) sound level in the block[e] | Distance from the expected level | $\|x - \mu\|$ | 15 dB | 19 dB | >1 | 5 dB | 1 dB | <1 | 5 dB | 1 dB | <1 |
| Precision-weighted mean | Expected sound level and its precision, $\pi = 1/\sigma^2$[e] | Precision-weighted distance from the expected level | $\pi \cdot \|x - \mu\|$ | 0.18 $dB^{-1}$ | 1.00 $dB^{-1}$ | >1 | 0.06 $dB^{-1}$ | 0.05 $dB^{-1}$ | <1 | 0.06 $dB^{-1}$ | 0.05 $dB^{-1}$ | <1 |
| No predictive model | Adaptation to the stimulus history within the chunk[f] | — | — | — | — | ~1 | — | — | ~1 | — | — | ~1 |

| Observed data | Deviant Gain 95% HDI | 1st standard Gain 95% HDI | 3rd standard Gain 95% HDI |
|---|---|---|---|
| facilitated profiles | [0.89, 1.12] | [0.76, 1.10] | [0.85, 1.12] |
| suppressed profiles | [0.93, 1.08] | [0.84, 1.05] | [0.87, 1.06] |

**Table S1. Predicted gain of the deviant and standard responses under different predictive models of the sequence.**

All predictions assume that prediction-error increases monotonically with the quantity in 'Quantity setting the prediction error' column.

[a] Variable block

[b] Stereotyped block

[c] The 3rd position sound differs from the first two in 6 of 7 chunks in the variable block and in 1 of 5 chunks in the stereotyped block. A deviant conforms to the learned pattern in the variable block but violates it in the stereotyped block.

[d] The 1st standard in the chunk has no preceding sound within the chunk, so the model makes no prediction for it.

[e] Values are for a 75 dB SPL standard and a 55 dB SPL deviant: $\mu$ = 70 and 74 dB SPL, and $\sigma^2$ = 83.3 and 19.0 $dB^2$, in the variable and stereotyped blocks respectively. The other tested configurations (75 dB SPL standard with a 45 dB SPL deviant; 45 dB SPL standard with a 75 dB SPL deviant) give the same directions. If $\mu$ and $\sigma^2$ are computed separately for each position, the deviant prediction is unchanged and the first standard shows no change.

[f] A model expecting that each sound repeats the preceding sound within the chunk, not learned from the block, makes the same predictions.
